# Rapamycin Treatment Reduces Brain Pericyte Constriction in Ischemic Stroke

**DOI:** 10.1101/2024.03.18.584333

**Authors:** Daniel J Beard, Lachlan S Brown, Gary P Morris, Yvonne Couch, Bryan A Adriaanse, Christina Simoglou Karali, Anna M Schneider, David W Howells, Zoran B Redzic, Brad A Sutherland, Alastair M Buchan

**Affiliations:** Acute Stroke Programme, Radcliffe Department of Medicine, University of Oxford, Oxford, UK; School of Biomedical Sciences and Pharmacy, University of Newcastle, Newcastle, Australia; Tasmanian School of Medicine, College of Health and Medicine, University of Tasmania, Hobart, Australia; Nuffield Department of Clinical Neurosciences, University of Oxford, Oxford, UK; Department of Oncology, University of Oxford, Oxford, UK; Department of Physiology, Faculty of Medicine, Kuwait University, Kuwait City, Kuwait

**Author notes:** Corresponding authors: Dr Daniel Beard, Assoc. Prof. Brad Sutherland. denotes co-first and co-senior authorship (Daniel J Beard, Lachlan S Brown, Brad A Sutherland, Alastair M Buchan). **Competing Interests** AMB is a senior medical science advisor and co-founder of Brainomix, a company that develops electronic ASPECTS (e-ASPECTS), an automated method to evaluate ASPECTS in stroke patients. All other authors declare no conflict of interest.

**Keywords:** Brain Pericytes, Experimental Stroke, Rapamycin, mTOR, RhoA, Calcium

## Abstract

The contraction and subsequent death of brain pericytes may play a role in microvascular no-reflow following the re-opening of an occluded artery during ischemic stroke. Mammalian target of rapamycin (mTOR) inhibition has been shown to reduce motility/contractility of various cancer cell lines and reduce neuronal cell death in stroke. However, the effects of mTOR inhibition on brain pericyte contraction and death during ischemia have not yet been investigated. Cultured pericytes exposed to simulated ischemia for 12 hours *in vitro* contracted after less than 1 h, which was about 7h prior to cell death. Rapamycin significantly reduced the rate of pericyte contraction during ischemia, however, it did not have a significant effect on pericyte viability at any time point. Rapamycin appeared to reduce pericyte contraction through a RhoA-dependent pathway, independent of changes in intracellular calcium. Using a mouse model of middle cerebral artery occlusion, rapamycin significantly increased the diameter of capillaries underneath pericytes and increased the number of open capillaries 30 minutes following recanalization. Our findings suggest rapamycin may be a useful adjuvant therapeutic to reduce pericyte contraction and improve cerebral reperfusion post-stroke.

## Introduction

Pericytes are vascular mural cells located between endothelial cells and astrocyte end-feet within the basal lamina of capillaries, and are found in particularly high density on capillaries of the central nervous system compared to other organs [1]. Recent evidence suggests that pericytes are contractile and regulating the luminal size of capillaries both *in vitro* and *in vivo* [2, 3]. This is particularly relevant in ischemic stroke as irreversible pericyte constriction (“death in rigor”) has been implicated as a major component of the “no-reflow” in preclinical models, a phenomenon where microvascular patency and blood flow are not restored even after large artery occlusions are removed [4]. This reduces the effectiveness of stroke therapies solely focusing on revascularization such as thrombolysis and thrombectomy in model systems [2, 5]. The no-reflow phenomenon also occurs in human stroke with one third of stroke patients reported to have microvascular perfusion deficits despite successful arterial recanalization [6]. Preventing brain pericyte death in rigor is therefore a potential therapeutic target to improve microvascular perfusion post-recanalization [7].

The Stroke Treatment Academic Round Table X (STAIR X) guidelines recommend prioritizing cytoprotective approaches that exert pleiotropic effects on multiple targets of the ischemic cascade and protect all brain components affected in stroke [8, 9]. Amplifying the brain’s intrinsic neuroprotective pathways is one such multimodal approach for developing new treatments for stroke. Mammalian target of rapamycin (mTOR) has been shown to play an important role in cell death after stroke [10]. In states of sufficient energy supply, mTORC1 is activated, signaling anabolic cellular processes such as protein synthesis, cell proliferation, cytoskeletal formation, and cellular contractility/motility [10]. Conversely, low cellular energy, i.e., increased AMP:ATP ratio, such as in cerebral ischemia, induces Tuberous Sclerosis-1 (TSC1, also known as Hamartin) and TSC2 (Tuberin) to form the TSC complex, subsequently down-regulating mTORC1 [11, 12], which has been shown to be protective in models of brain ischaemia [10]. Therefore, enhancement of endogenous mTOR inhibition, and hence, augmentation of cell-preserving effects, is an appealing target for brain cytoprotection in cerebral ischemia.

Rapamycin (Sirolimus™) [13], an immunosuppressant frequently used in clinical practice [14], inhibits mTORC1. Inhibition of mTORC1 with rapamycin has been shown to reduce infarct volume and decrease neurological deficits in animal models of cerebral ischemia [15]. Rapamycin can also protect multiple cells within the brain including neurons [16], and brain endothelial cells [17] in *in vitro* models of ischemia [18]. Rapamycin treatment has also been shown to improve collateral blood flow in experimental stroke by increasing endothelial nitric oxide synthase activity [19]. Further, rapamycin can decrease Rho-A mediated cytoskeletal reorganization and thus motility/contractility in different cancer cell lines [20]. As rapamycin is already in clinical use with a known side effect profile, potential extrapolation to the stroke clinic is a distinct possibility. In the current study, we investigated if rapamycin could reduce pericyte contractility both *in vitro* and following experimental stroke *in vivo* and investigated the mechanism by which rapamycin may exert this effect.

## Materials and Methods

### Cell Culture

Primary cultures of pericytes from rat brain were gown using a previously described method for extracting primary brain endothelial cells (BECs) from two-to three-month-old Wistar rats [5], and modified to enable pericyte growth [21]. Human Brain Vascular Pericytes (HBVP) (#1200) were purchased from ScienCell Research Laboratories, cultured in complete pericyte medium, and used between passages 6 and 8.

### Immunocytochemistry

Immunocytochemistry for canonical pericyte markers was performed on primary rat pericytes to confirm the purity of cultures. We have previously characterised HBVPs extensively [22-24].

### Oxygen glucose deprivation (OGD)

Rat brain pericytes were exposed to either OGD or normoxia (control) conditions for 2-12 hours.

### Western Blotting

Western blotting was used to investigate changes in phosphorylated and total ribosomal protein S6 (t-S6 and p-S6, downstream indicators of mTORC1 activity) in rat pericytes following exposure to 2 or 8 h of OGD or normoxia and treatments with vehicle, 10 nM rapamycin or 100 nM rapamycin.

### Measurement of rat pericyte contractility

Pericyte contractility was measured using an electrical impedance system that detected changes in the contact area between rat pericytes and the culture dish, as described previously by Neuhaus et al. [24] during exposure to 12 h of OGD or normoxia and treatment with vehicle, 10 nM rapamycin, 100 nM rapamycin or 10nM rapamycin plus 1μM U46619, a PGH_2_ analogue that is a potent and stable thromboxane A_2_ (TP) receptor agonist that activates RhoA [25, 26].

### Measurement of rat pericyte cell death

Flow cytometry analysis of Annexin V–APC and Propidium Iodide (PI), was used to measure apoptosis/necrosis in rat pericytes after exposure to 2, 8 or 12 h of OGD or normoxia and treatment with vehicle, 10 nM rapamycin or 100 nM rapamycin.

### Human pericyte contraction and calcium imaging during chemical ischemia

We utilised chemical ischaemia (hereinafter referred to as ischaemia) with antimycin-A and sodium iodoacetate to block oxidative phosphorylation and glycolysis, to study the effect of 30nM of rapamycin on human pericyte contraction and Ca^2+^ flux during ischemia.

### Middle Cerebral Artery Occlusion (MCAO) and Cerebral Blood Flow Measurement

The intraluminal filament MCAO model was performed for 60 minutes in adult 3–4-month-old male NG2-DsRed mice as previously described [27]. At the commencement of MCAO, mice were randomised to treatment with an IP injection of either 1 mg/kg rapamycin or saline (vehicle control).

### Tissue fixation, processing, and imaging

After reperfusion, mice were terminally anaesthetised and transcardially perfused with PFA and FITC-Albumin + gelatin to obtain a microvascular cast of perfused vessels, as described previously [28]. Sections were labelled with Isolectin GS-IB4-Alexa Fluor 647 conjugate to detect blood vessels and DAPI to detect cell nuclei. Images were processed in ImageJ and analysed blinded to treatment group and ROI using independently developed ImageJ macro scripts.

### Statistical Analysis

Detailed methodological descriptions can be found in the Supplementary Materials.

## Results

### Rapamycin reduced mTORC1 activity in rat pericytes during normoxia and oxygen glucose deprivation (OGD)

Initially, expression of pericyte markers PDGFRβ and desmin in cultured rat brain pericytes were confirmed by immunocytochemistry (Supplementary Fig. 1). To confirm successful mTORC1 inhibition with rapamycin treatment, we measured the phosphorylation of the downstream target of mTOR, S6 ribosomal protein [29]. Both 10nM and 100nM doses of rapamycin significantly reduced the p-S6/t-S6 ratio in pericytes after 2 h of Normoxia (Fig. 1a and b; 10nM: 0.1 ± 0.04, 100nM: 0.08 ± 0.03 versus Vehicle: 1.0 ± 0.2, both p < 0.0001). Only 100nM of rapamycin significantly reduced the p-S6/ t-S6 ratio in pericytes after 8 h of Normoxia (Fig. 1c and d; 10nM: 0.2 ± 0.009, 100nM: 0.0 ± 0.0 versus Vehicle: 1.0 ± 0.6, 10nM: p = 0.3, 100nM: p = 0.01).

**Fig 1.**
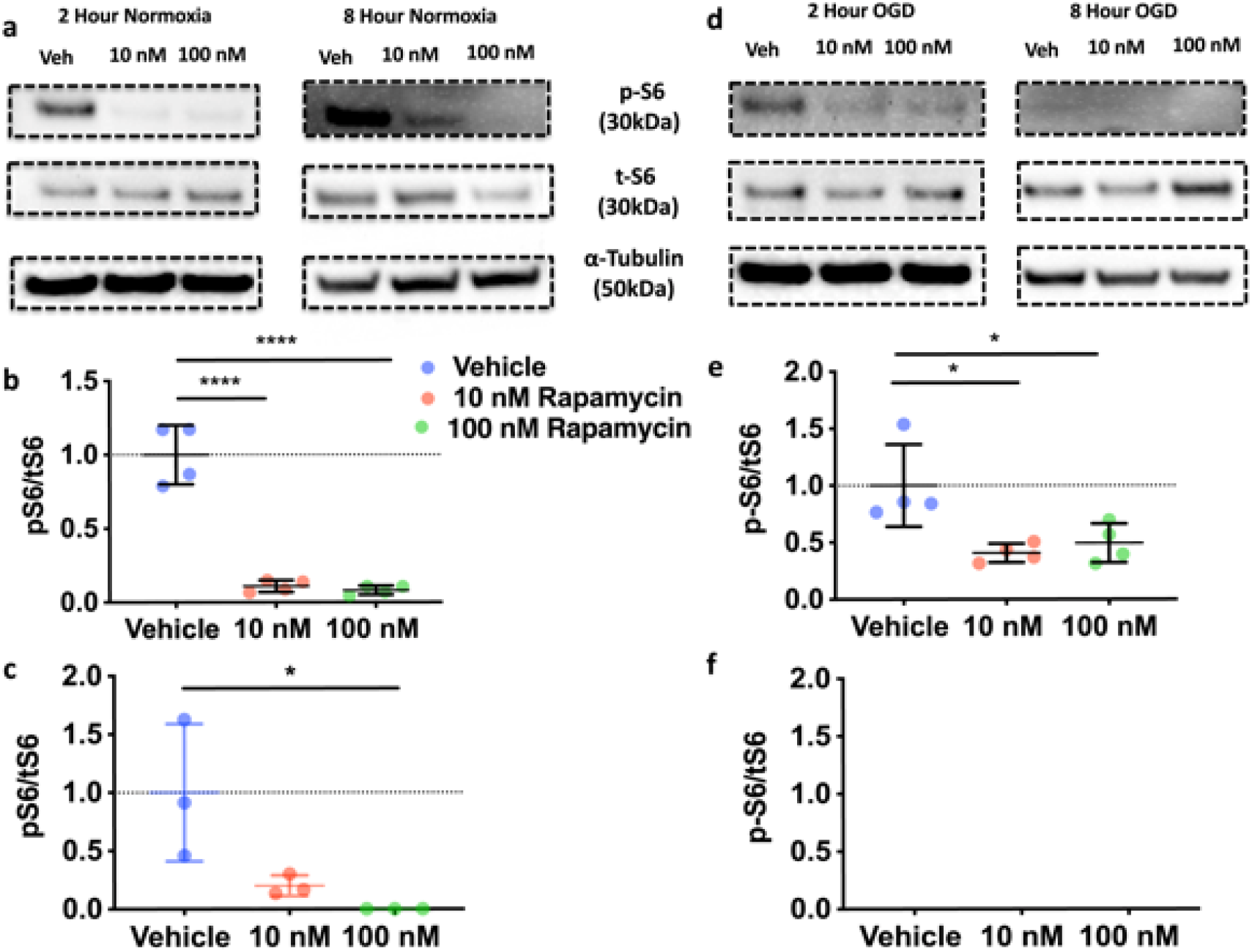
Effect of rapamycin treatment on mTORC1 activity. (**a**) Representative blots for pS6, t-S6 and loading control, tubulin after 2 h and 8 h of Normoxia. (**b**) Quantification of p-S6 relative to t-S6; (indicating mTOR activity) after 2 h normoxia. One way ANOVA (Treatment, F(2,9) = 76.26, p < 0.0001) with Dunnett’s test multiple comparisons test to compare treatment groups. (**c**) Quantification of p-S6 relative to t-S6 after 8 h normoxia. Kruskal Wallis: H (2) = 7.448, p = 0.0036 with Dunn’s multiple comparisons test to compare treatment groups. (**d**) Representative blots after 2 h and 8 h of OGD. (**e**) Quantification of p-S6 relative to t-S6 after 2 h OGD. Kruskal Wallis (Treatment, H (2) = 7.538, p = 0.0107) with Dunn’s multiple comparisons test to compare treatment groups. (**f**) Quantification of p-S6 relative to t-S6 after 8 h OGD. Each value was mean ± SD from 3-4 samples (split over 3-4 gels/blots). * p < 0.05, **** p < 0.0001.

We then tested whether rapamycin could inhibit mTORC1 in OGD. Both 10nM and 100nM of rapamycin significantly reduced p-S6/t-S6 ratio in pericytes after 2 h of OGD (Fig. 1d and e; 10nM: 0.407 ± 0.08233, 100nM: 0.4935 ± 0.17 versus Vehicle: 1.00 ± 0.3609, 10nM: p = 0.01, 100nM: p = 0.03). There was no detectable p-S6 in any of the groups during 8 h of OGD, however, t-S6 was still detected (Fig. 1d and f). Full membrane scans of blots can be found in Supplementary Fig. 2-7.

### Rapamycin reduced rat pericyte contractility without affecting pericyte cell death

We analyzed the effects of ischemia and rapamycin treatment on pericyte contraction using an electrical impedance system (iCelligence) [24]. Exposure of rat pericytes to 12 h of OGD resulted in a decline in cell index (indicative of contraction) (Fig. 2a). This contraction was due to the ischemia rather than the media change, as the pericyte cell index did not decline after changing media to normoxia media (Supplementary Fig. 8). We tested the effects of rapamycin treatment on the slope of the cell index curve (indicating the rate of pericyte contraction). There was no significant effect of either dose of rapamycin on the slope of the curve in the first hour of OGD (Fig. 2b; 10nM: -0.05 ± 0.01, 100nM: -0.05 ± 0.03 versus Vehicle: -0.06 ± 0.01, 10nM: p = 0.9, 100nM: p = 0.7). However, both doses of rapamycin significantly reduced the slope of the cell index curve between 1-2 h of OGD (Fig. 2c; 10nM: -0.01 ± 0.03, 100nM: -0.01 ± 0.03 versus Vehicle: -0.04 ± 0.03, 10nM: p = 0.03, p = 0.03). Only 100nM rapamycin significantly reduced the slop of the cell index between 2-3 h of OGD (Fig. 2d; 10nM: -0.04 ± 0.04, 100nM: -0.03 ± 0.03 versus Vehicle: -0.08 ± 0.06, 10nM: p = 0.1, 100nM: p = 0.04). There was no effect of rapamycin treatment on the slope of the cell index between 3-4 h of OGD (Fig. 2e; 10nM: -0.03 ± 0.03, 100nM: -0.04 ± 0.04 versus Vehicle: -0.05 ± 0.03, 10nM: p = 0.3, 100nM: p = 0.5).

**Fig 2.**
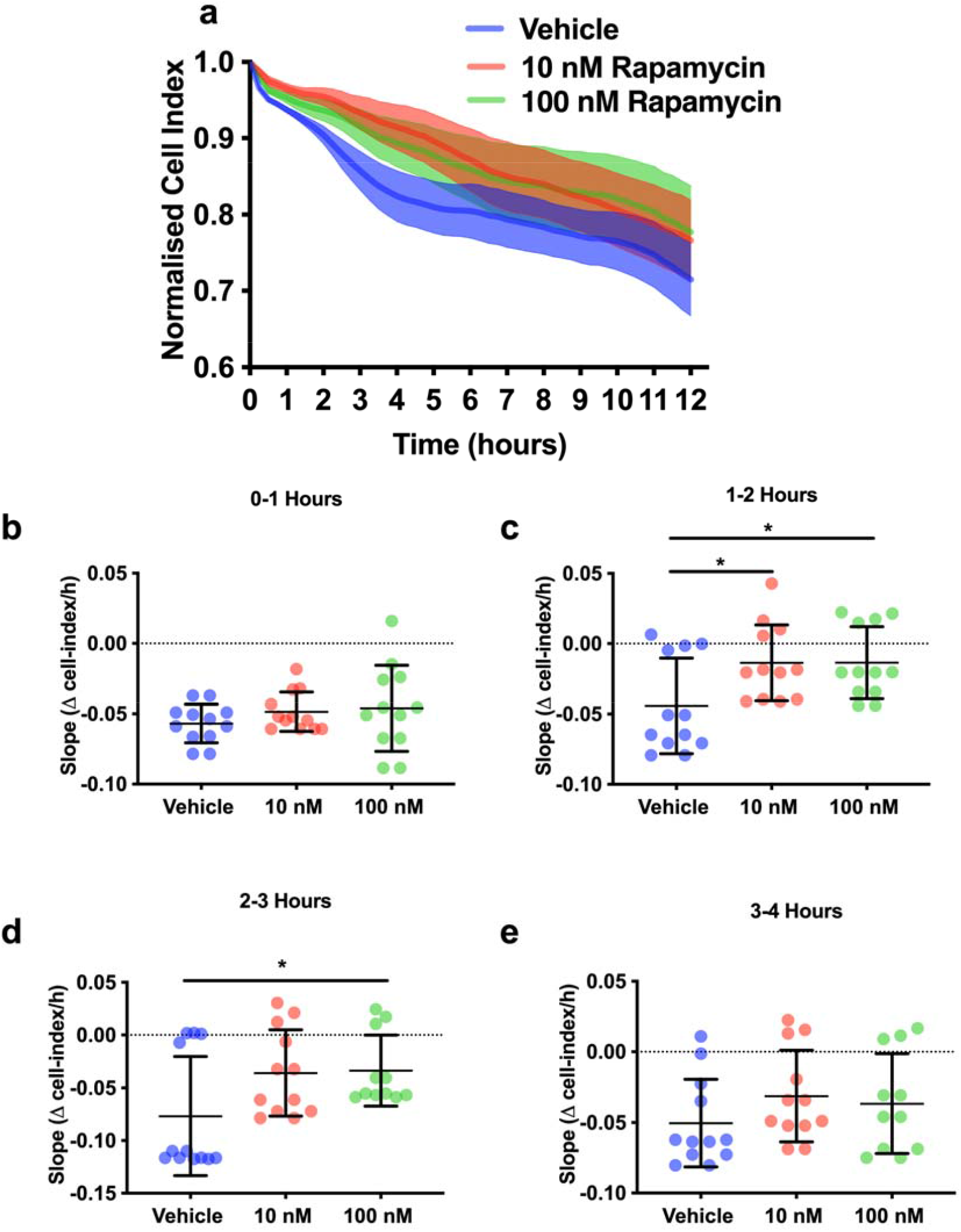
Rapamycin reduces pericyte contractility during OGD. (**a**) Average normalized cell index for vehicle (blue), 10nM rapamycin (red) and 100nM rapamycin (green) treated cells during 12 h of OGD. (**b**) Slope of cell index 0-1 h post-OGD. Kruskal Wallis (Treatment, H (2) = 1.054, p = 0.5904) with Dunn’s multiple comparisons test to compare treatment groups. (**c**) Slope 1-2 h post-OGD. One way ANOVA (Treatment, F(2,33) = 4.428, p = 0.0198) with Dunnett’s test multiple comparisons test to compare treatment groups. (**d**) Slope 2-3 h post-OGD. Kruskal-Wallis (Treatment, H(2) = 6.34, p = 0.0420) with Dunn’s multiple comparisons test to compare treatment groups. (**e**) Slope 3-4 h post-OGD. One way ANOVA (Treatment, F(2,32) = 1.74, p = 0.357) with Dunnett’s test multiple comparisons test to compare treatment groups. Each value was mean ± SD from 4 wells per treatment group from 3 independent cultures. * p < 0.05.

We performed flow cytometry for propidium iodide and annexin-V (see Supplementary Figure 9 for example heat maps), to confirm that the observed effect of rapamycin was due to its effect on contraction, rather than a consequence of changes in cell death and/or detachment. There was no reduction in cell viability at 2 h of OGD (Fig. 3a; OGD Vehicle: 88.9 ± 6.0 % versus Normoxia Vehicle: 88.9 ± 5.2 %), suggesting that the reduction in cell index reflects pericytes contraction rather than cell death and the subsequent detachment. Furthermore, rapamycin had no effect on pericyte viability at 2 h in the OGD group (Fig. 3a; 10nM: 86.4 ± 5.9 %, 100nM: 84.3 ± 5.9 % versus Vehicle: 88.9 ± 6.0 %, 10nM: p = 0.6, 100nM p = 0.2). No significant pericyte death was recorded until 8 and 12 h of OGD (Fig. 3b-e; 12h OGD Vehicle: 38.9 ± 16.1 % versus Normoxia Vehicle: 80.4 ± 8.5 %). Rapamycin had no effect on pericyte viability in the OGD group at 12 hours (Fig. 3b; 10nM: 38.9 ± 17.5 %, 100nM: 40.2 ± 18.7 % versus Vehicle: 38.9 ± 16.1 %, 10nM: p = 0.98 100nM p = 0.96).

**Fig 3.**
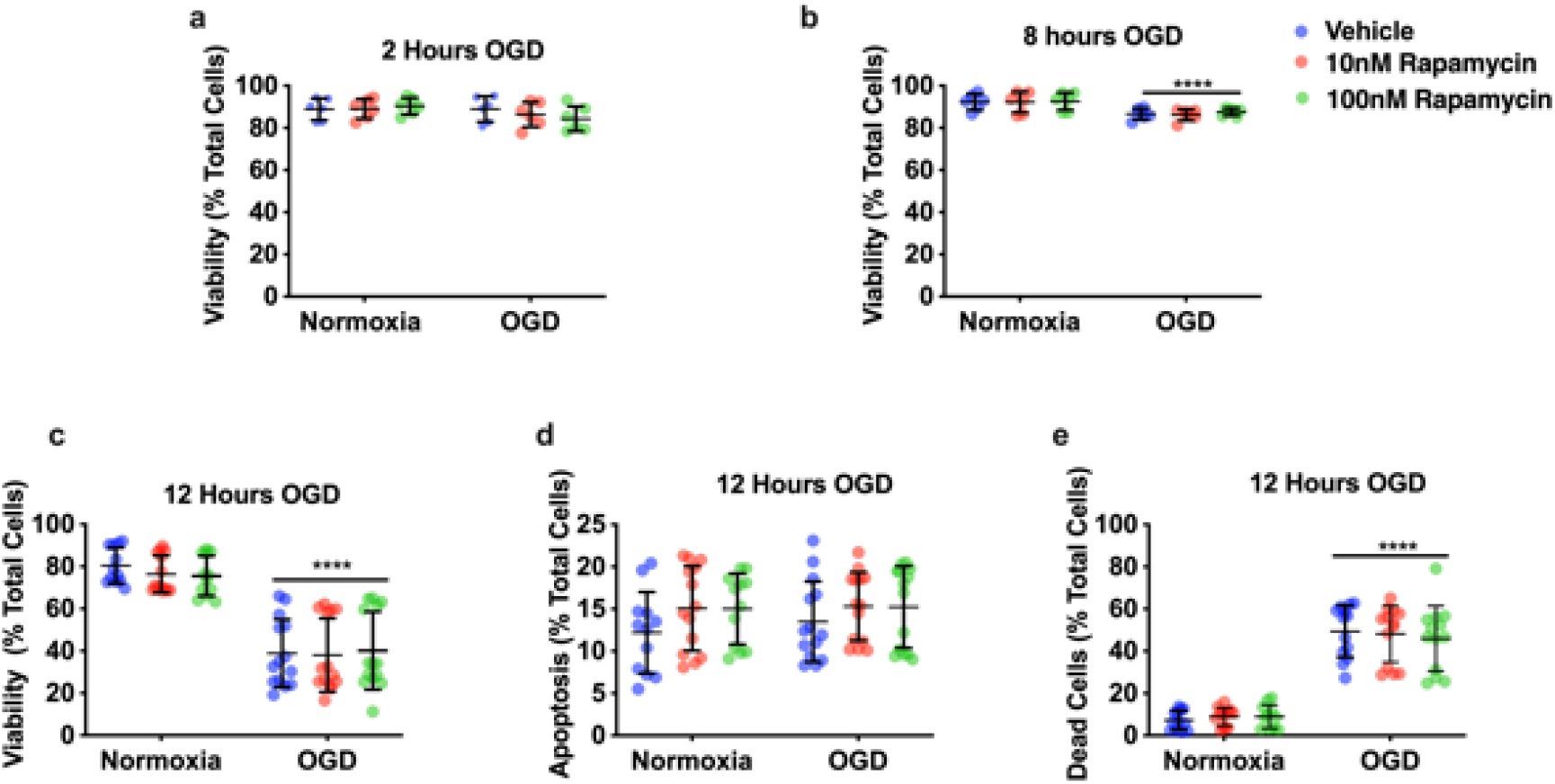
Rapamycin did not affect pericyte cell death during OGD. (**a**) Pericyte viability after 2 h OGD (% of cells negative for Annexin-V (AV) and Propidium Iodide (PI) staining). Two-way ANOVA, OGD, F (1,30) = 2.632, p = 0.3799; Treatment, F (2.30) = 0.2771, p = 0.7599. (**b**) Pericyte viability after 8 h OGD. Two-way ANOVA, OGD, F (1,30) = 22.64, p < 0.0001; Treatment, F (2.30) = 0.21, p = 0.8118. (**c**) Pericyte viability after 12 h OGD. Two-way ANOVA, OGD, F (1,74) = 149, p < 0.0001; Treatment, F (2, 74) = 0.2121, p = 0.8094. (**d**) Apoptosis of pericytes after 12 h OGD (% cells positive for AV but negative for PI). Two-way ANOVA, OGD, F (1,74) = 0.313, p = 0.5743; Treatment, F (2, 74) = 2.169, p = 0.1216. (**e**) Pericyte cell death 12 h after OGD (% cells positive for PI +/- Annexin V). Two-way ANOVA, OGD, F (1,69) = 264.4, p < 0.0001; Treatment, F (2, 69) = 0.07593, p = 0.9270. 2 and 8 h OGD: each value was mean ± SD from 2 wells per treatment group from 3 independent cultures with 2 wells per group. 12 h OGD: each value was mean ± SD from 4 wells per treatment group from 3 independent cultures. **** p < 0.0001 for effect of OGD.

### Rapamycin prevented human pericyte contractility to ischemia but did not inhibit calcium flux

We wanted to determine if calcium levels were associated with pericyte contraction in response to ischemia [2, 24], and whether rapamycin could modulate this effect. For these experiments, we used human brain vascular pericytes (HBVPs) since they were used in our previous work [24]. We tested the effects of ischemia and rapamycin treatment on the net area under the curve (nAUC), an indicator of the extent of pericyte cell membrane retraction (contraction), in a single cell imaging assay we have used previously [22]. Exposure of human pericytes to ischemia resulted in a retraction of the cell membrane at 20 min of ischemia (Fig. 4a, c) and a corresponding significant reduction in normalised nAUC over time compared to control treatment (Fig. 4d; Ischaemia + Vehicle: -1.4 ± 1.8 versus Vehicle: 0.4 ± 1.9, p < 0.0001). Rapamycin significantly reduced pericyte membrane contraction (Fig. 4b, c) and significantly increased nAUC compared to ischemia alone (Fig. 4d; Ischaemia + Rapamycin: -0.3 ± 1.9 versus Ischaemia + Vehicle: -1.4 ± 1.8, p = 0.03). We then assessed whether rapamycin affected Ca^2+^ changes in pericytes in response to ischemia using the calcium indicator Fura-2. Ischemia resulted in no change in Ca^2+^ flux compared to vehicle alone (Ischaemia: 1.0 ± 0.6 versus Vehicle: 0.89 ± 1.3, p = 0.8). Rapamycin caused a small but significant increase in calcium flux during ischemia compared to ischemia alone (Fig. 4e, f; Rapamycin + Ischemia: 1.5 ± 0.5 versus Ischemia: 1.0 ± 0.6, p = 0.01). These results suggest that rapamycin can prevent ischemia-induced pericyte contraction downstream of calcium entry.

**Fig 4.**
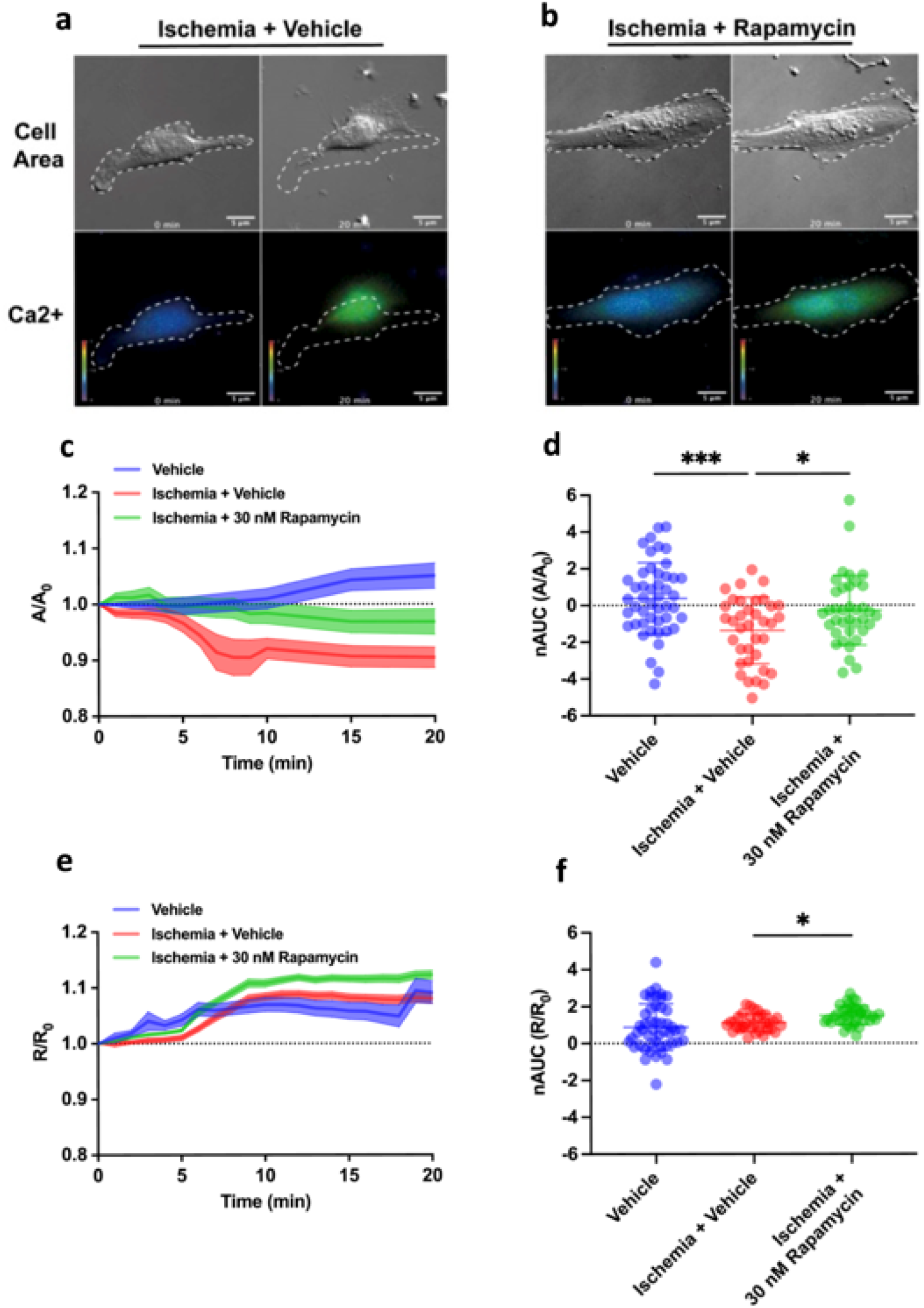
Rapamycin prevents pericyte contraction in response to chemical ischemia. Representative DIC and fluorescent fura-2 AM Ca^2+^ flux images of pericytes treated with: (**a**) chemical ischemia + vehicle (DMSO) or (**b**) rapamycin, at baseline (0 min) and 20 min time points. The white dotted outline indicates cell membrane boundaries for each representative cell at baseline. Colorimetric scale represents intracellular Ca^2+^ levels with the ratiometric Fura2-AM Ca^2+^ indicator. Lower right scale bar = 10μm. For each cell: (**c**) normalised cell area (A/A_0_) and (**e**). normalised Fura2-AM calcium flux (R/R_0_), compared to baseline was calculated over time following each treatment. The dotted line represents baseline cell area or calcium flux. The dark lines represent the mean and the shaded area represents the SEM for each treatment group. Number of individual cells analysed for each group: Vehicle (DMSO) (N = 47); Ischemia (N = 36); Ischemia + Rap (Rapamycin) (N = 37); over 6 replicate cultures. (**d**) Net area under the curve (nAUC) of A/A_0_ and (**f**). R/R_0_ was calculated for each treatment group. Bars represent mean ± SD. One-way ANOVA (Treatment, F (2,117) = 8.785, p =0.0003) with Dunnett’s multiple comparisons test was used to compare A/A_0_ groups. Kruskal-Wallis test (Treatment, H (2) = 15.28, p = 0.0005) with Dunn’s multiple comparisons test was used to compare R/R_0_ groups. * p < 0.05, **** p < 0.0001.

### Activation of RhoA blocks the effect of rapamycin on rat pericyte contractility during OGD

Since rapamycin reduced pericyte contractility by acting downstream of calcium entry, we hypothesised that rapamycin may be altering the sensitivity of the cytoskeleton to calcium, specifically through the RhoA myosin light chain kinase pathway [20]. To determine the role of RhoA in the effect of rapamycin on rat pericyte contraction, we repeated the rapamycin OGD iCelligence experiments in the presence of U46619, a PGH_2_ analogue that is a potent and stable thromboxane A_2_ (TP) receptor agonist that activates RhoA. Exposure of rat pericytes to OGD resulted in an initial increase in cell index over the first hour of OGD followed by a decline in cell index over the following 11 h of exposure (Fig. 5a). Rapamycin + U46619 significantly reduced the positive slope of the cell index relative to vehicle in the first hour of OGD (Fig. 5b; Rapamycin + U46619: 0.02 ± 0.006 versus Vehicle: 0.03 ± 0.25, p = 0.03). There was no significant difference in negative slope of the cell index between vehicle and rapamycin + U46619 at 1-2 h (Fig. 5c; Rapamycin + U46619: -0.13 ± 0.07 versus Vehicle: -0.2 ± 0.09, p = 0.1), 2-3 h (Fig. 5d; -0.2 ± 0.1 versus Vehicle: -0.2 ± 0.2, p = 0.97) and 3-4 h (Fig. 5e; Rapamycin + U46619: -0.2 ± 0.1 versus Vehicle: -0.2 ± 0. 1.0) of OGD.

**Fig 5.**
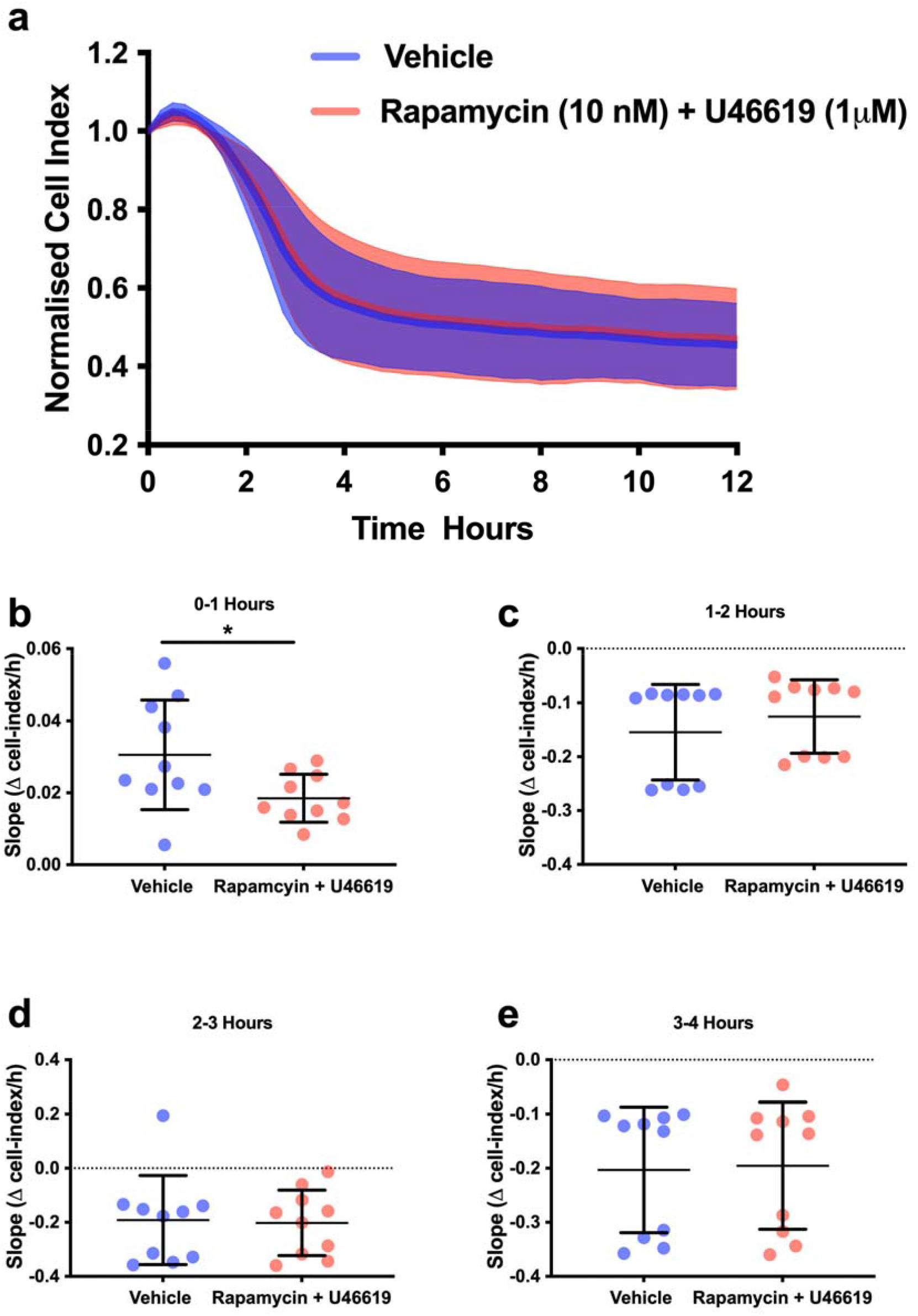
RhoA activation blocks reduced pericyte contractility by rapamycin during OGD. (**a**) Average normalized cell index for vehicle (blue) and 10nM rapamycin + 2μM U46619 (red) treated cells during 12 h of OGD. (**b**) Slope of cell index between 0 and 1 h post-OGD. Unpaired t-test: t (18) = 2.3, p = 0.0336. (**c**) Slope between 1 and 2 h. Mann-Whitney U test: U = 29, p = 0.1230. (**d**) Slope between 2 and 3 h. Mann-Whitney U test: U = 49, p = 0.9705. (**e**) Slope between 3 and 4 h. Mann-Whitney U test: U = 49, p = 0.9705. Each value was mean ± SD from 5 samples from 2 independent cultures. * p < 0.05.

### Rapamycin increases vessel width and the number of open vessels in the striatum after MCAO

Given that rapamycin can delay ischemia-induced contraction in pericytes (Fig. 2 and 4), we next investigated whether rapamycin could affect vessel width under pericytes following MCAO as a measure of pericyte contraction. Laser Doppler Flowmetry monitoring of blood flow in the upper layers of the somatosensory cortex did not reveal any change in the extent of reperfused blood flow with rapamycin treatment (Supplementary Fig. 10). However, Laser Doppler Flowmetry can only measure overall flow coming from all vessels in a small volume of tissue directly below the probe. We therefore performed a microvascular luminal cast using FITC-albumin perfusion to analyse changes in individual vessel perfused luminal width beneath pericyte soma after treatment with rapamycin [28]. The area of perfusion deficit during ischemia in this model encompasses the MCA territory which includes the striatum and the cerebral cortex containing the somatosensory cortex, these regions were chosen for analysis (Fig. 6a). Despite 30 min of recanalisation of the MCA following 60 min MCAO, there appeared to be areas devoid of FITC-albumin luminal labelling, suggestive that some capillaries were closed (Fig. 6b). Interestingly, many of these areas with closed vessels (vessels devoid of luminal labelling) appeared to be associated with pericytes (Fig. 6c).

**Fig 6.**
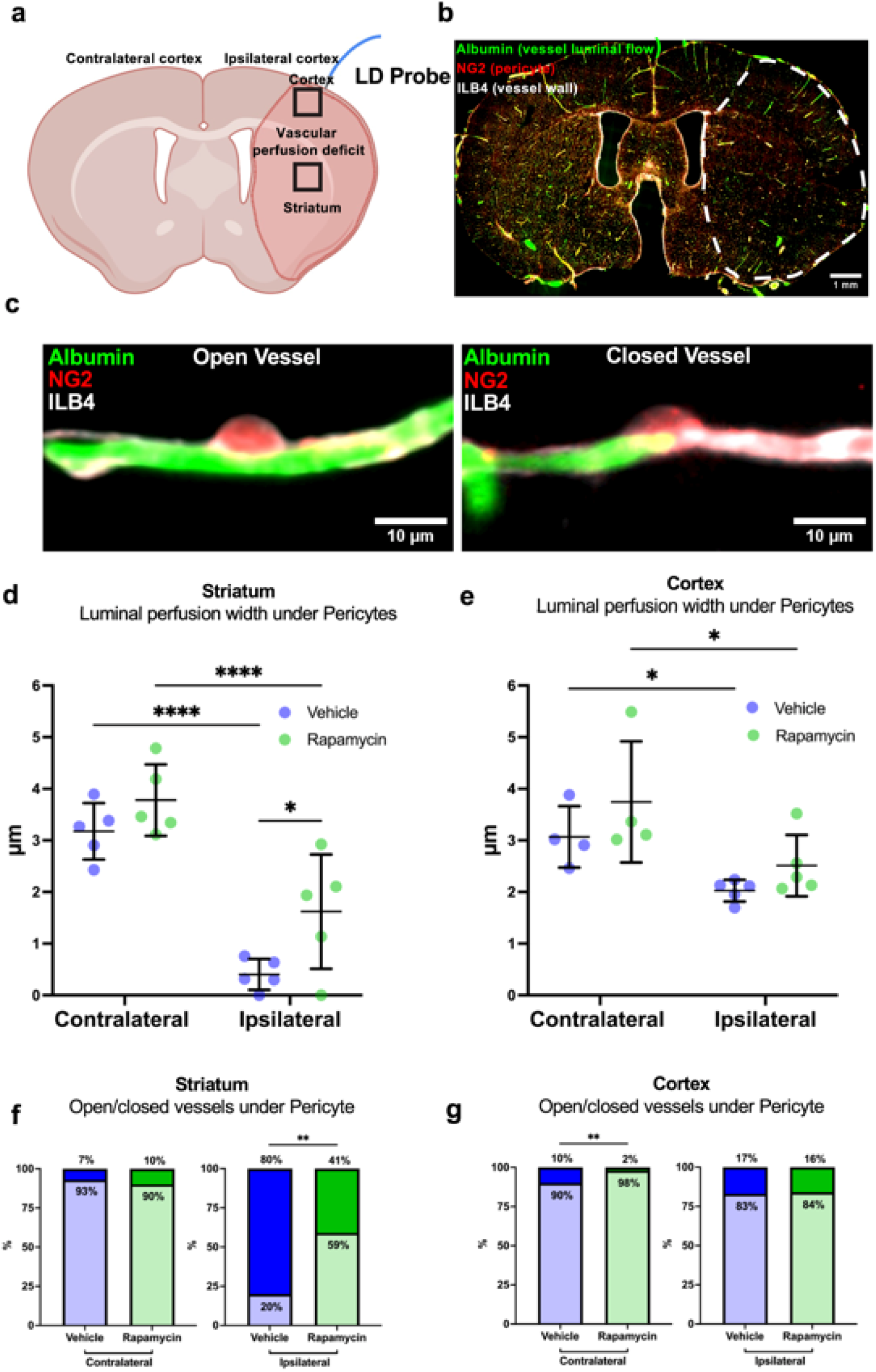
Rapamycin increases luminal width and the proportion of open vessels in the striatum following MCAO. (**a**) Schematic of mouse brain coronal section under bregma demonstrating contralateral and ipsilateral hemispheres, relative location of laser doppler (LD) probe, the area of vascular perfusion deficit in the ipsilateral hemisphere, and regions of interest sampled. (**b**) Example fluorescent image of whole coronal brain section labelled with FITC-albumin (green), NG2DsRed (Red), and isolectin B4 (white), with the area of vascular perfusion deficit annotated with white dotted line. (**c**) Example images of capillaries with luminal FITC-albumin labelling (left) and reduced luminal FITC-albumin labelling (right) under pericyte soma. (**d**,**e**). Quantification of mean perfused vessel luminal width under pericytes in striatum and cortex in both the contralateral and ipsilateral hemispheres following 60 min MCAO and 30 min reperfusion. Individual data points are presented with mean ± SD and represent mean data determined across N = 1086 individual vessel width measurements from N = 10 animals across both treatment groups. Repeated measures 2way ANOVA [(**d**) Hemisphere, F (1,8) = 162.9, p < 0.0001; Treatment, F (1,8) = 4.831, p = 0.0592, and (**e**) Hemisphere, F (1,5) = 38.93, p < 0.0015; Treatment, F (1,9) = 2.961, p = 0.1194] with Sidak’s multiple comparisons test was used to compare treatment and hemisphere groups. When values were missing, a mixed-effects model with Sidak’s multiple comparisons test was used. (**f**,**g**). Vessel data (from **d**,**e**) were binned based on luminal width being 0 (closed) or >0 (open) in striatum and cortex. Data are presented as a percentage of open or closed vessels. Fisher’s exact test was used to compare groups. * p < 0.05, ** p < 0.01, **** p < 0.0001.

Analysis of luminal width underneath pericyte soma revealed that MCAO had caused a substantial decrease in vessel width in the ipsilateral striatum compared to the corresponding contralateral hemisphere (Fig. 6d; Ipsilateral Vehicle: 0.4 ± 0.3 μm versus Contralateral Vehicle: 3.2 ± 0.5 μm, p < 0.0001). Rapamycin significantly increased luminal width in the ipsilateral striatum compared to vehicle (Fig. 6d; Rapamycin: 1.6 ± 1.1μm versus Vehicle: 0.4 ± 0.3μm, p= 0.03). Rapamycin did not affect luminal width in the contralateral striatum compared to vehicle (Fig. 6d; Rapamycin: 3.8 ± 0.7 μm versus Vehicle: 3.2 ± 0.5 μm, p = 0.4). There was a significant decrease in vessel width in the ipsilateral cortex compared to the corresponding contralateral hemisphere (Fig. 6e, Ipsilateral Vehicle: 2.0 ± 0.2 μm versus Contralateral Vehicle: 3.1 ± 0.6 μm, p = 0.0111). However, rapamycin did not significantly alter vessel width compared to vehicle in the ipsilateral (Rapamycin: 2.5 ± 0.6μm versus Vehicle: 2.0 ± 0.2 μm, p = 0.2) or contralateral (Rapamycin: 3.7 ± 1.2 μm versus Vehicle: 3.1 ± 0.6 μm, p = 0.3) hemispheres (Fig. 6e).

It was notable that some vessels had no FITC-albumin signal indicative of closed vessels both in the striatum and cortex following MCAO. We therefore quantified the proportion of open (presence of some FITC-albumin signal under pericytes) and closed (zero FITC-albumin signal under pericytes) vessels. Consistent with the vessel width data, the ipsilateral striatum only had 20% of capillaries open while the contralateral striatum had 90% of capillaries open. Rapamycin significantly increased percentage of open capillaries compared to vehicle (Fig. 6f; Rapamycin: 59% versus Vehicle: 20%, p = 0.0059). Rapamycin did not significantly alter the percentage of open capillaries in the ipsilateral cortex (Fig. 6g; Rapamycin: 84% versus Vehicle: 83%, p = 0.9). Interestingly, rapamycin significantly increased the percentage of open capillaries in the contralateral cortex compared to vehicle (Fig. 6g; Rapamycin: 98% versus Vehicle: 90%, p = 0.002).

### MCAO induces pericyte loss which is not prevented by rapamycin

Our *in vitro* data had shown that rapamycin does not reduce pericyte death in response to OGD (Fig. 4), and so we determined whether rapamycin could prevent pericyte loss following MCAO. We found that pericyte number, as defined by the presence of NG2DsRed signal on vessels, was reduced in the ipsilateral compared to the contralateral striatum (Fig. 7a, b; Ipsilateral Vehicle: 22 ± 15 pericytes/mm^2^ versus Contralateral Vehicle: 95 ± 27 pericytes/mm^2^, p < 0.0001). Rapamycin did not significantly affect pericyte number in the striatum of either the ipsilateral (Rapamycin: 19 ± 18 pericytes/mm^2^ versus Vehicle: 22 ± 15 pericytes/mm^2^, p = 0.99) or contralateral (Rapamycin: 86 ± 30 pericytes/mm^2^ versus Vehicle: 95 ± 27 pericytes/mm^2^, p = 0.8) hemispheres (Fig. 7b). There was no significant change in pericyte number in the ipsilateral cortex compared to the contralateral cortex (Fig. 7c; Ipsilateral Vehicle: 75 ± 29 pericytes/mm^2^ versus Contralateral Vehicle: 94 ± 26 pericytes/mm^2^, p = 0.6). Rapamycin had no significant effect on pericyte numbers in the cortex of either the ipsilateral (Rapamycin: 66 ± 37 pericytes/mm^2^ versus Vehicle: 75 ± 29 pericytes/mm^2^, p = 0.9) or contralateral (Rapamycin: 90 ± 23 pericytes/mm^2^ versus Vehicle: 94 ± 26 pericytes/mm^2^, p = 0.99) hemispheres (Fig. 7c).

**Fig 7.**
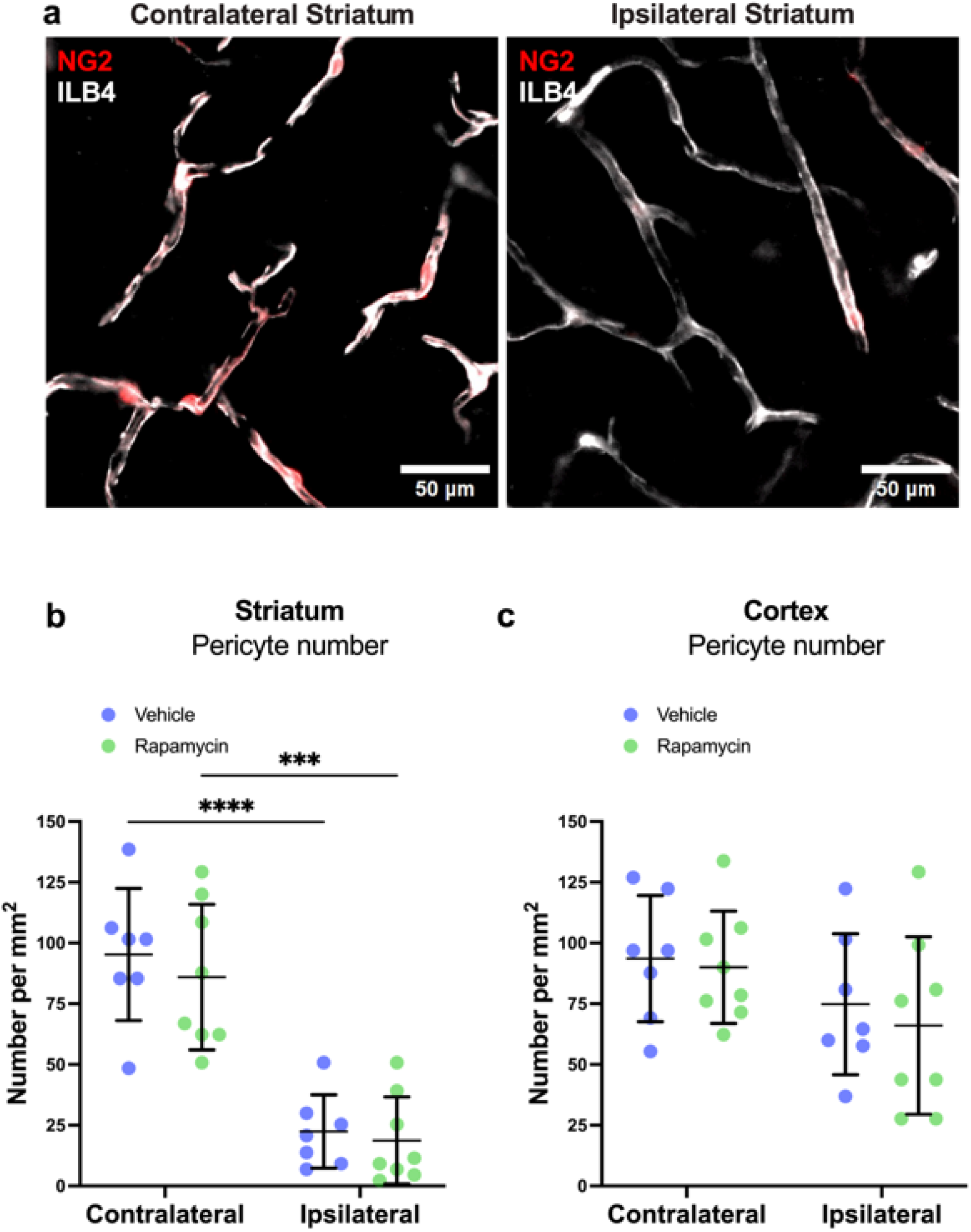
Rapamycin had no effect on the reduction in pericyte number in the striatum following MCAO. (**a**) Example images of capillaries (ILB4, white) with pericytes (NG2, red) in contralateral (left) and ipsilateral (right) striatum of mice subject to 60 minutes MCAO followed by 30 minutes reperfusion. (**b**,**c**) Quantification of pericyte number in striatum and cortex. Individual data points represent individual mice overlayed with mean ± SD. 2-way ANOVA [(**b**) Hemisphere, F (1,13) = 70.43, p < 0.0001; Treatment, F (1,8) = 0.5430, p = 0.4743, and (**c**) Hemisphere, F (1,13) = 4.579, p = 0.0519; Treatment, F (1,13) = 0.2980, p = 0.5944] with Tukey’s multiple comparisons test was used to compare groups. **** p < 0.0001.

## Discussion

This is one of few reports of a potential therapeutic intervention aimed to reduce pericyte contraction in response to ischemia. Rapamycin reduced cultured pericyte contraction during ischemia *in vitro* (using both OGD and ischemia paradigms). We demonstrate that rapamycin exerts its anti-contractile effect downstream of intracellular calcium changes, which involved reduction of signalling through the RhoA GTPase pathway. Furthermore, these effects were independent of changes in pericyte cell death. Finally, we highlight the translational potential of rapamycin by showing that it increases capillary diameter at pericyte locations and the number of open capillaries in mice following MCAO.

Our results, along with previous *in vitro* investigations, indicate that pericytes contract before they die. We demonstrated that pericytes begin to contract within an hour of OGD onset but do not begin to die until at least 7 h later. These results are in line with our laboratory’s previously published work, which showed that HBVPs exposed to ischemia begin to contract within an hour after treatment onset but do not die until 24 h later [24]. Hall *et al*. [2] showed that pericytes in cortical slices begin to contract 15 min after the onset of ischemia and die in significant numbers after 1h. Hall *et al*. [2] reported that pericytes die at 24 h after 90 min of MCAO, but the timeframe of pericyte contraction was not measured in their study. Meanwhile, Yemisci *et al*. [5] reported pericyte constriction of capillaries at 6 h of reperfusion following 2 h MCAO but did not report pericyte cell death *in vivo*. Here we showed that MCAO significantly reduced capillary diameter at pericyte locations and decreased the number of open vessels in the ipsilateral striatum 30 minutes following recanalisation. These findings are broadly consistent with those of Yemisci *et al*. [5]. Furthermore, a recent study by Qiu et al. using two-photon imaging found that there was significant vasoconstriction in the areas of vessels covered by pericyte somas as well as processes at 3 h post-recanalization [30]. A very recent study by Shrouder et al. [4] showed that 87% of pericytes constrict during 60 minutes of intraluminal MCAO ischemia and remain constricted following reperfusion in mice. This constriction was significantly associated with capillary stalls (stalled red blood cells in a capillary) at pericyte soma during ischemia and out to 24 hours of reperfusion. These findings demonstrate that pericytes present a promising therapeutic target to counteract no-reflow in stroke.

Rapamycin treatment significantly slowed pericyte contraction during OGD and increased capillary diameter at pericyte soma and increasing the number of open capillaries at pericyte soma following MCAO. This strongly suggests that rapamycin is reducing the constriction of pericytes both *in vitro* and *in vivo*. This is the first study to report that rapamycin can reduce pericyte contraction. One previous study managed to influence capillary diameter by using nanoparticles that contained adenosine conjugated to the lipid squalene (which allowed a prolonged circulation of this nucleoside) in mice that were subjected to MCAO / reperfusion. Among other observed neuroprotective effects, the adenosine nanoparticles could prevent capillary clogging, which strongly suggests a therapeutic effect on cells of the neurovascular unit, however, the role of pericytes in these effects on capillaries was not specifically investigated [31].

Rapamycin’s anti-contractile effect does not appear to be mediated by changes in pericyte cell death *in vitro* as pericytes did not die until approximately 12 h of OGD, and rapamycin had no effect on cell death at any time point investigated. This is in line with previous studies showing that cultured brain pericytes undergo rapid cell cycle arrest [32], but do not die during 4 h of OGD [21, 32]. Previous studies investigating other cell types showed rapamycin can reduce cell death in cultured neurons [16] and cultured brain endothelial cells [17] during OGD. Furthermore, rapamycin has been shown to reduce astrocyte proliferation, migration and production of inflammatory mediators to OGD *in vitro* [18], reduce blood-brain barrier permeability and reduce infarct volume in animal models of stroke [13, 15]. The lack of pericyte protection (at least *in vitro*) afforded by rapamycin may be explained by the hypothesis that OGD itself down-regulates mTORC1 in vehicle-treated cells, which means that by the time the pericytes start to die between 8 and 12 h, there would be no additional mTORC1 inhibition effected by rapamycin treatment.

Although we did not directly assess pericyte cell death *in vivo*, we did observe a significant reduction in NG2DsRed signal following 60 minutes of MCAO and 30 minutes of ischemia. This could be suggestive of early pericyte cell injury leading to pericyte membrane shedding or leakage of the cytoplasm through damaged membranes. Shrouder et al. [4] also reported a loss of EGFP signal from pericytes at 90 minutes following reperfusion, however they did not directly investigate pericyte cell death using TUNEL staining until 24 hours, which revealed an increase in pericyte death at this timepoint. Therefore, it remains to be determined if pericyte cell death occurs as early as 30-and 90-minutes following reperfusion. We did, however show there was no significant difference in the reduction of NG2DsRed signal between vehicle and rapamycin treatment groups. This could suggest that rapamycin has little influence on pericyte damage in the early phases of reperfusion following stroke.

We showed that rapamycin reduced the contraction of human pericytes in response to ischemia without inhibiting influx of intracellular calcium. Previous studies showed that pericyte contraction during CI in brain slices was dependent on rapid extracellular calcium entry through voltage-gated calcium channels (VGCCs) [2] and that derivatives of rapamycin can reduce the opening of VGCCs in hippocampal neurons [33], while rapamycin itself did not exert activity at VGCCs [34]. In this study, we found only a small increase in calcium levels between ischaemia alone and ischaemia plus rapamycin at the time of maximal effect of rapamycin on contraction. This suggests that rapamycin could be influencing pericyte contractility downstream of calcium entry/release.

Activation of RhoA with the thromboxane-A2 agonist U46619 reduced the effect of rapamycin on pericyte contraction during OGD. RhoA has been shown to regulate the contractility of bovine retinal pericytes via myosin light chain kinase and alpha-smooth muscle actin [35] and to induce brain pericyte contraction in cortical slices and *in vivo* [36, 37]. Whereas the two other Rho GTPases, Rac and Cdc42, primarily alter cell shape via changes in the cortical actin cytoskeleton leading to localised changes in the shape of the cell membrane (not contraction *per se*) [38]. Rapamycin has also been shown to reduce RhoA expression and activation via mTORC1 inhibition and subsequent reduction in S6 kinase and eukaryotic translation initiation factor 4E-binding protein 1 (4EBP1) activity to reduce cytoskeletal rearrangement in human rhabdomyosarcoma and Ewing sarcoma cell lines [20]. Our findings strongly suggest that rapamycin may reduce contractility by reducing RhoA activity leading to a decrease in phosphorylation of the myosin light chain, thus reducing the sensitivity of the contractile apparatus of pericytes to calcium (Supplementary Fig. 11). Shrouder et al. [4] demonstrated that topical application of the ROCK-inhibitor fasudil over the stroke cortex could reverse pericyte constriction and capillary stalls during 70 min MCAO and at 90 minutes and 24 hours of reperfusion in mice. This is in line with our findings that pericyte constriction can be reversed and further highlights the important role of the Rho-A/ROCK pathway in inducing pericyte constriction post-stroke. Therefore, blocking this pathway with rapamycin or fasudil could become therapeutic strategies to reverse no-reflow after stroke.

In conclusion, we found that rapamycin reduces the rate of pericyte contraction during OGD and ischemia, and the reduction in contraction in OGD was mediated by a RhoA-dependent mechanism rather than a reduction in cell death or inhibiting intracellular calcium. Complementing our *in vitro* findings, we found that rapamycin also increased the diameter of capillaries at pericyte soma and increased the number of open capillaries in the striatum post-recanalization in mice. This strongly suggests that rapamycin also reduces pericyte constriction following stroke *in vivo*. These findings have important therapeutic relevance for preventing no-reflow after successful re-opening of occluded arteries after ischemic stroke.

## Acknowledgements

We would also like to thank Dr. Helen Ferry for her assistance in setting up the flow cytometry protocols used in this manuscript.

## Declarations

### Ethical Approval

All animal procedures were approved by the University of Tasmania Animal Ethics Committee (A0016160 and A0018608) and were compliant with the Australian NHMRC Code of Practice for the Care and Use of Animals for Scientific Purposes.

### Funding

DJB, YC, AMS, BAS and AMB were funded by the Medical Research Council UK (MR/M022757/1). DJB was funded by National Health and Medical Research Council Australia (APP1182153). AMB, and AMS funded by Leducq Foundation for Cardiovascular and Neurovascular Research (Consortium International pour la Recherche Circadienne sur l’AVC). AMB is an Einstein Visiting Fellow (EVF□2021-619), funded by the Einstein Foundation, Berlin. BAS and LSB were funded by the Rebecca L. Cooper Foundation and National Health and Medical Research Council Australia (APP1137776, APP2003351). CSK was funded by Cancer Research UK.

### Availability of data and materials

The data that support the findings of this study are available from the authors upon reasonable request.

