## Supplementary information for "Rapamycin Treatment Reduces Brain Pericyte Constriction in Ischemic Stroke"

Translational Stroke Research

Daniel J Beard^1,2#*^, Lachlan S Brown^3#^, Gary P Morris^3^, Yvonne Couch^1^, Bryan A Adriaanse^4^, Christina Simoglou Karali^5^, Anna M Schneider^1^, David W Howells^3^, Zoran B Redzic^6^, Brad A Sutherland^3#*^, Alastair M Buchan^1#^

^1^ Acute Stroke Programme, Radcliffe Department of Medicine, University of Oxford, Oxford, UK

^2^ School of Biomedical Sciences and Pharmacy, University of Newcastle, Newcastle, Australia

^3^ Tasmanian School of Medicine, College of Health and Medicine, University of Tasmania, Hobart, Australia

^4^ Nuffield Department of Clinical Neurosciences, University of Oxford, UK

^5^ Department of Oncology, University of Oxford, UK

^6^ Department of Physiology, Faculty of Medicine, Kuwait University, Kuwait

* Corresponding authors:

Dr Daniel Beard

\# denotes co-first and co-senior authorship

Supplementary Information

**Supplementary methods**

***Cell Culture***

Briefly, pericytes were cultured under standard incubation conditions (37^°^C; 5% CO2 in 95% humidified air) as described earlier [1], but without addition of Puromycin. Thus, both BECs and pericytes initially grew in cell culture medium that consisted of DMEM, 20% FBS, 1% penicillin/streptomycin, 0.1mg/mL ascorbic acid for 10 days with a full media change on Day 1, followed by a half media change every second day (P0 culture). Ten days after plating, pericytes largely overgrew BECs, at which point cells were passaged to uncoated, poly-L-lysine (Sigma Aldrich) treated 6 well plates and allowed to grow to ~70% confluence before being used for experimental procedures (P1 culture). This step eliminated BECs from cultures, as they could not produce enough basal lamina components and thus, do not attach to uncoated plastic [2].

***Immunocytochemistry***

Rat pericytes were seeded at 30,000 cells per well in 8 well Millicell® EZ Slides (Millipore) and allowed to adhere. After fixation (4% paraformaldehyde for 10 min), autofluorescence quenching (1% sodium borohydride for 10 min) and blocking (10% donkey serum for 1 h), the cultured cells were incubated overnight at 4^°^C in 1% serum/PBS that contained the following primary antibodies: cluster of differentiation 31 (CD31; goat polyclonal; R&D Systems; 1:100), beta-type platelet-derived growth factor receptor (PDGFRβ; goat polyclonal; R&D Systems; 1:100) and desmin (rabbit polyclonal; AbCam; 1:100). After washing, the cultures were incubated for 2 h at room temperature in the dark with secondary antibodies (anti-goat and anti-rabbit Alexa488-conjugated; AbCam; 1:400). Cytoskeletal actin was counterstained using Alexa594-conjugated phalloidin (Sigma-Aldrich; 1:500) and nuclei were counterstained with DAPI (Sigma-Aldrich). Images were acquired using a Zeiss Axioscope 2 (Carl Zeiss) with appropriate filter settings and acquisition parameters.

***Oxygen Glucose Deprivation***

Rat brain pericytes were exposed to either OGD or normoxia (control) conditions for 2-12 hours. For control experiments, rat brain pericyte cultures were incubated in glucose- and pyruvate-free DMEM (Gibco #11966025) that contained a reduced serum concentration (2%) and was supplemented with 5 mM glucose and 1.25 mM pyruvate in 5% CO2 in humidified room air at 37°C (control media). For OGD experiments, a hypoxic glove box (Coy Laboratories) maintained with 0% O_2_, 5% CO_2_ in N_2_ at 37°C was used. All media and buffers were left in the glove box for at least 12 h prior to OGD experiments. When transferred to the glove box, cell cultures were incubated in normoxia media, without supplementation of glucose- and pyruvate (OGD media). The same control and OGD conditions were used for pericyte contractility, western blotting and cell death analysis.

***Measurement of pericyte contractility***

We used an electrical impedance system to detect changes in the contact area between rat pericytes and the culture dish, as described previously by Neuhaus et al. [3]. In brief, rat pericytes were plated on impedance plates (E-Plate L8 PET; ACEA Biosciences) and allowed to proliferate in the iCelligence platform (ACEA Biosciences) for 48 h prior to testing. Pericytes were then exposed to OGD for 12 h and treated with vehicle (0.005% ethanol in media), 10 nM rapamycin (Sigma #553210) or 100 nM rapamycin. The same vehicle and drug concentrations were used for all subsequent experiments. To determine the role of RhoA in the effect of rapamycin on pericyte contraction, we treated pericytes with vehicle (ethanol + 0.35% DMSO in media) or 10 nM rapamycin + 1 μM U46619 (Cayman Chemicals #16450), a thromboxane A2 agonist that activates RhoA [4, 5]. 10 nM rapamycin was selected for these experiments as there was no additional benefit of 100 nM rapamycin on reducing pericyte contractility in initial experiments. Data are presented as normalized cell index (a unitless parameter, automatically derived from recorded impedance values in iCelligence software and normalized to the last time point pre-OGD), with the rate of contraction calculated by the slope of the cell index.

***Measurement of rat pericyte cell death***

Flow cytometry was used to measure apoptosis/necrosis in rat pericytes after exposure to 2, 8 or 12 h of OGD or normoxia as well as treatments with vehicle, 10 nM rapamycin or 100 nM rapamycin. At the end of the exposure period, cells were stained with Annexin V–APC and Propidium Iodide (PI), according to the manufacturer’s instruction (Biolegend) (for details see the Supplementary Material). Flow cytometry was carried out on a BD LSRII Fortesa (BD Biosciences) with optimised parameters (Supplementary Fig. 1). Data were collected using BD FACSDiva, Version 8.0 and analysed with FlowJo X, Version 10.0.7r2 (FlowJo).

***Western Blotting***

Cells were lysed in cell lysis buffer through cell scraping, incubated at 4°C for 30 min and centrifuged at 4°C for 30 min at 11,700 g. Supernatant was collected for Western blotting. Protein concentration (mg/mL) of cell lysates were determined (RD DC Protein Assay, Biorad using a DC (Biorad, UK). 10 μg of sample per well was run on NuPAGE 4-12% Bis-Tris gels (Invitrogen) and then transferred to nitrocellulose using an iBlot2 dry transfer system (Invitrogen). Membranes were blocked in 5% BSA for 1 h at room temperature. Membranes were immunoblotted with primary antibodies against phospho-S6 at Ser235/236 (p-S6; rabbit monoclonal; Cell Signaling Technology; 1:2000), total S6 protein (t-S6; rabbit monoclonal; Cell Signalling Technology; 1:8000), and α-Tubulin (mouse monoclonal; Abcam; 1:4000) diluted in 5% BSA 0.1% TX-100 and incubated overnight at 4^o^C. Secondary antibodies (HRP-conjugated, goat anti-rabbit or goat anti-mouse immunoglobulin, 1:1000, Dako) were applied to the membranes for 1 h at room temperature. The immunoblots were visualised and quantified on a Biorad ChemiDoc^TM^ MP imaging system (Biorad) using ECL-advanced detection reagents (Invitrogen). The density of the bands was measured using Biorad Image Lab software v6.0.1 (Biorad). The loading controls were performed by analysis of the t-S6 protein and α-tubulin protein. S6 phosphorylation at Ser235/236 (an index of S6 kinase activity and readout of upstream mTORC1 activity) was expressed as the ratio of S6 phosphorylation to t-S6 protein, to account for variability in t-S6 protein between samples.

***Human pericyte contraction and calcium imaging during chemical ischemia***

For all live single cell imaging experiments, primary HBVP were cultured on glass coverslips and imaged on a Nikon Ti Live Cell Microscope and imaged using the same methodology as described previously [6], with modification. Experiments with HBVP were performed between passages 6-8.HBVP were incubated with Fura 2-AM ratiometric calcium indicator (1 μM, Life Technologies) diluted in imaging buffer [1 X HBSS (no Ca2+, no Mg^2+^, no phenol red), 15 mM HEPES, 30 mM glucose, 1 mM MgCl_2_, 2 mM CaCl_2_ in Milli-Q H2O, pH 7.4] and maintained at 37°C for 10 min. HBVP were imaged using differential interference contrast (DIC) microscopy on a temperature-controlled 37°C upright Nikon Ti Live Cell Microscope with a 40x oil immersion objective and EMCCD camera (Photometrics Evolve 512x512). Pericytes were only imaged provided there were no other interfering cells in the field of view. For contractility assessment, following baseline images, imaging buffer was removed and replaced with imaging buffer containing vehicle (0.1% v/v DMSO), chemical ischemia solution (5 μM Antimycin A, Sigma #A8674; 500 μM Sodium iodoacetate, Sigma; concentrations determined from [3]), and rapamycin (30 nM, Adooq Bioscience). Live DIC images (to visualise cell area) and fluorescent (340 nm and 380 nm excitation and dual 510 nm emission; to visualise Ca^2+^ flux) images were recorded at 1 min intervals. 7-12 cells were imaged per coverslip within the same session with an automated stage facilitating repeated return to their XY coordinates for recording over time. Images were processed with NIS Elements Analysis software and exported to ImageJ for further analysis. To determine the extent of change relative to baseline over a period of time, normalised cell area and normalised Ca^2+^ flux was calculated as described in previous literature [6].

***Middle Cerebral Artery Occlusion and Cerebral Blood Flow Measurement***

Adult 3–4-month-old male NG2-DsRed mice (Jackson Laboratories #008241) were used for all MCAO experiments. All animal procedures were approved by the University of Tasmania Animal Ethics Committee (A0016160 and A0018608) and were compliant with the Australian NHMRC Code of Practice for the Care and Use of Animals for Scientific Purposes. Mice were housed in standard conditions with ad libitum access to food and water and on a 12 h light:dark cycle (light phase was 7:00 AM-7:00 PM). The intraluminal filament MCAO model (Longa method) was performed as previously described [7]. Mice were anaesthetised with isoflurane at 5% in O2 delivered at 1 L/min in a box for induction, and 2-4% in O_2_ delivered at 0.5 L/min intranasally through a nose cone for maintenance (Advanced Anaesthesia Specialists, Australia). A silicone-coated filament (602145PK10; 20-25 g mouse, 602345PK10; 25-35 g mouse; Doccol Corporation, USA) was advanced into the right external carotid artery and up the internal carotid artery to occlude the origin of the right MCA. At the commencement of MCAO, mice were treated with an IP injection of either 1 mg/kg rapamycin (Sapphire Bioscience #A10782-10MM-D) or saline (vehicle control). After 60 min, the filament was retracted to allow recanalisation of the MCA and reperfusion of the brain. To measure cerebral blood flow (CBF) during the MCAO and reperfusion periods, a laser Doppler probe (moorVMS-LDF1, Moor Instruments) was positioned in a silicon probe holder (PHDO, Moor Instruments) perpendicular to the right temporal skull at 3 mm lateral and 1.5 mm caudal to bregma. Occlusion was confirmed by a sudden drop in CBF to <30% of baseline.

***Blinding, animal numbers, and exclusions***

Animals were randomly assigned to treatment and surgical groups (rapamycin vs vehicle (saline); MCAO vs sham) and researchers were blinded to treatment groups by preparing treatments in identical vials and assigning treatments to arbitrary groups A and B. Animals were excluded if a blood flow drop to <30% of baseline during MCAO was not exhibited, they were not able to achieve a stable anesthetic plane, or a terminal bleed or haemorrhage occurred as a result of the surgery. Out of 28 animals that were entered into the study, there were 4 exclusions.

***Laser Doppler signal processing and analysis***

LD data were recorded at 100Hz using LabScribe (IWorks) and down sampled to 0.1Hz for export (bins of 10 second data) for each animal to determine changes in CBF during common carotid artery occlusion (CCAO, middle cerebral artery occlusion (MCAO), middle cerebral artery reperfusion (MCAR), and common carotid artery reperfusion (CCAR). Baseline traces were cut to a flat 5-minute baseline section prior to CCAO, and normalised to the mean over that baseline period. All CCAO periods were normalised to the last 5 minutes prior to MCAO. All MCAR periods were normalised to the first 2 minutes following MCAR.

***Tissue fixation and processing***

After 30 min of reperfusion, mice were terminally anaesthetised via IP injection of pentobarbital (300 mg/kg) and transcardially perfused at 9.6 mL/min with 37°C PBS + heparin (1 unit/mL; DBL Heparin Sodium Injection BP, Pfizer) for 1 min, then PBS + 4% w/v paraformaldehyde (Sigma) for 1 min, followed by PBS + 0.1% w/v FITC-Albumin (Sigma) + 5% w/v gelatin (Sigma) for 1 min to obtain a microvascular cast of perfused vessels, as described previously [8]. Mice were then incubated on ice for 30 min to set the gelatin, after which brains were removed, incubated in PBS + 4% PFA for 1 h at 4°C, washed 3 x in PBS and cryoprotected in PBS + 30% w/v sucrose (Sigma #S0389) for 24 h at 4°C. Brains were rapidly frozen by placing them in molds with Epredia Cryomatrix embedding resin (Thermofisher) on a floating foam tube holder on liquid nitrogen and then stored at -80°C. Brains were sectioned coronally at 40 μm with a CM1850 Cryostat (Leica Biosystems) at -18°C, and tissue slices were placed directly onto slides for analysis.

***Blood vessel labelling immunohistochemistry***

Slide-mounted tissue was permeabilised with 0.3% v/v Triton X-100 (Sigma) in Dako Antibody Diluent (Agilent) for 40 min at room temperature. Tissue was blocked using Dako Serum Free Protein Block (Agilent) for 60 min at room temperature. Isolectin GS-IB4- Alexa Fluor 647 conjugate (Thermofisher) was diluted 1:500 in Dako Antibody Diluent and incubated with the tissue for 24 h at 4°C in a humidified dark chamber. Sections were then washed 3 x 10 min in PBS, before being incubated in 1X Trueblack (Biotium) for 1 min to reduce the impact of autofluorescence, washed 3 x 2 min in PBS, rinsed in distilled water, then cover-slipped with Prolong Gold antifade reagent with DAPI (Thermofisher).

***Image acquisition and analysis***

Slides were imaged using a VS120 slide scanner (Olympus, Japan) as described previously [8]. After an overview (DAPI, Ex: 388 nm; Em: 448 nm; 2X magnification), tracing of tissue sections and focus-map generation, slides were scanned using Extended Focus Imaging (15 μm total range, 3 μm spacing) at 20X magnification in the DAPI (Ex: 388 nm; Em: 488 nm), FITC (Ex: 494 nm; Em: 530 nm), Texas red (Ex: 576 nm; Em: 625 nm), and Cy5 (Ex: 650 nm; Em: 670 nm) channels. Optimum exposure times were initially determined manually and then kept consistent for all images.

Images were processed in ImageJ and analysed blinded to treatment group using independently developed ImageJ macro scripts to allow semi-automated analysis and reduce image analysis bias (https://github.com/brownls23/NG2DsRed-IHC-analysis-.git). To detect NG2-DsRed positive capillary pericytes, binarised DAPI and NG2-DsRed soma-specific overlays were generated to calculate pericyte number, with a manual binary threshold matching step, DAPI-positive cell selection step, and removal of smooth muscle cell NG2DsRed signal step. FITC-Albumin specific overlays were generated with a manual binary threshold matching step and removal of large vessels. These images were used to analyse perfused vessel width underneath pericyte soma, with vascular association confirmed by positive ILB4 labelling. Width was defined as the minimum vascular width of FITC-albumin labelling beneath a NG2-DsRed positive pericyte soma and measured using the Straight-Line tool in ImageJ. Where one pericyte spanned two vessel branches, width of both vessels was measured. Where there was no FITC-Albumin signal present or FITC-Albumin signal stopped under a pericyte soma, perfused vessel width under pericyte was given a value of 0 μm. For open/closed analysis of perfused vessel width, width measurements were binned into two groups: 0 μm and >0 μm.

***Statistical Analysis***

Data were processed in Microsoft Excel and all statistical analysis was performed using GraphPad Prism 10 (GraphPad, USA). When data was tested for outliers, a ROUT test (Q = 1%) was used and outliers were removed from data sets unless they were biologically relevant. When comparing at least three independent variables, data were tested for normality of the residuals using the D'Agostino & Pearson test, or the Shapiro–Wilk test if n numbers were too small. When this data was normally distributed, variables were compared using Ordinary one-way ANOVA with Dunnett’s multiple comparisons test between groups. When this data was not normally distributed, variables were compared using Kruskal-Wallis test with Dunn’s multiple comparisons test between groups. When comparing how a response is affected by two factors where one of the factors was repeated, independence, normality, and sphericity was assumed, and variables were compared using Repeated measures (RM) two-way ANOVA with Sidak’s multiple comparisons test between groups. When comparing frequency of two categorical variables contingency table matrices were used, and variables compared using Fisher’s exact test. A p < 0.05 was considered statistically significant. Statistical tests used for each analysis are reported in figure legends. For ANOVAs, overall ANOVA results are reported in figure legends and results of *post hoc* tests are reported in the results text and are represented in each figure.

**Supplementary results**

***Cultures of rat pericytes express canonical pericyte markers***

Immunocytochemistry for canonical pericyte markers was performed on primary rat pericytes to confirm the purity of cultures. Staining with CD31 was negative indicating that there was no contamination of the primary culture with endothelial cells. PDGFRβ, a receptor known to be expressed by CNS pericytes [9], was found in high abundance in this cell population. All cells in the cultures also stained positive for desmin, a contractile cell marker for smooth muscle cells and pericytes [10] (Supplementary Fig. 1).

**Supplementary Figure**s

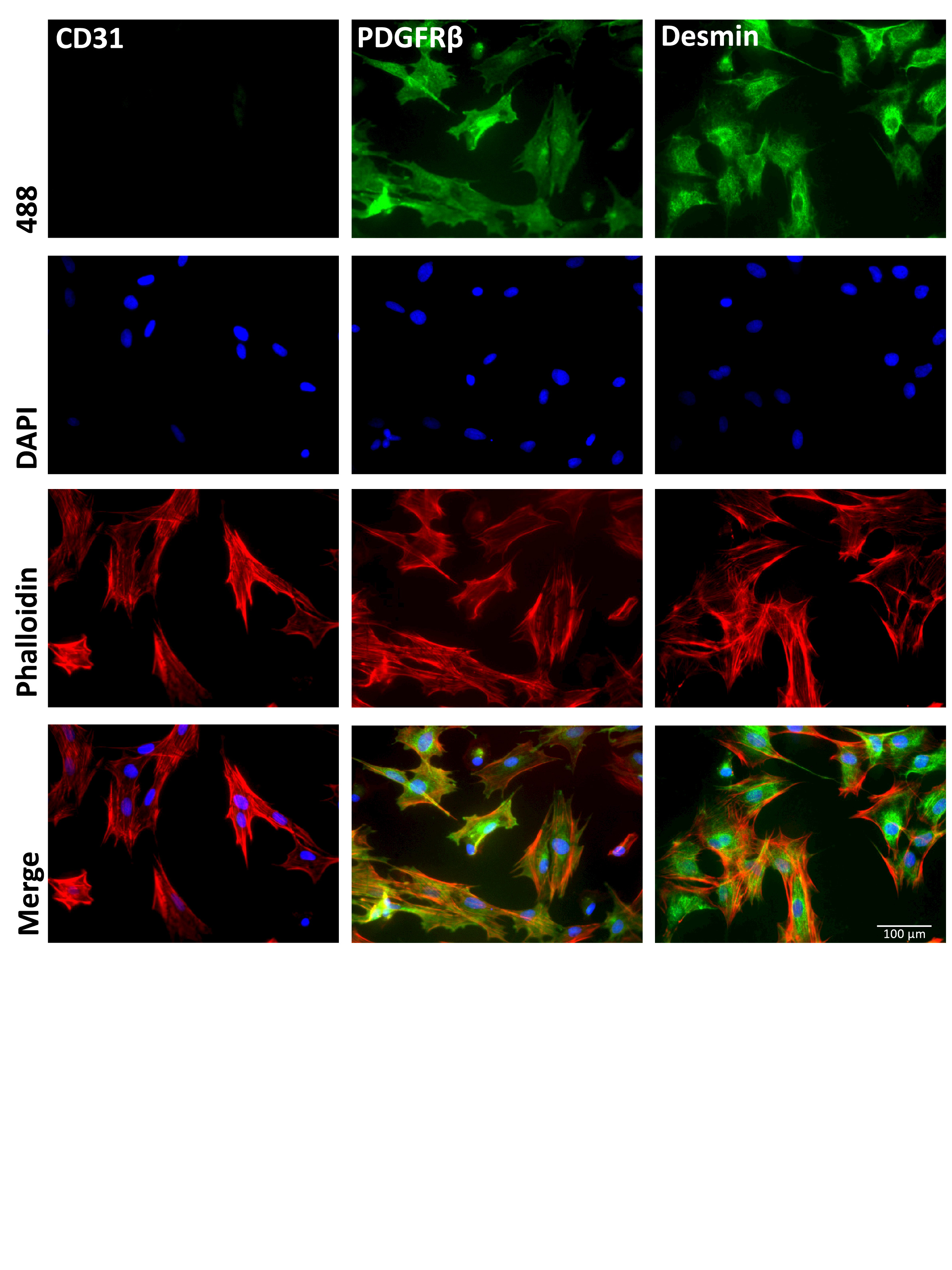

**Supplementary Fig 1** Immunocytochemistry of primary rat brain pericytes. Pericytes were fixed in 4% paraformaldehyde and stained for endothelial (CD31), and pericyte (PDGFRβ and Desmin) markers (all green – Alexa488) to confirm a pericyte phenotype. Cells were counterstained with a cytoskeletal marker (phalloidin – red) and a nuclear marker (DAPI – blue).

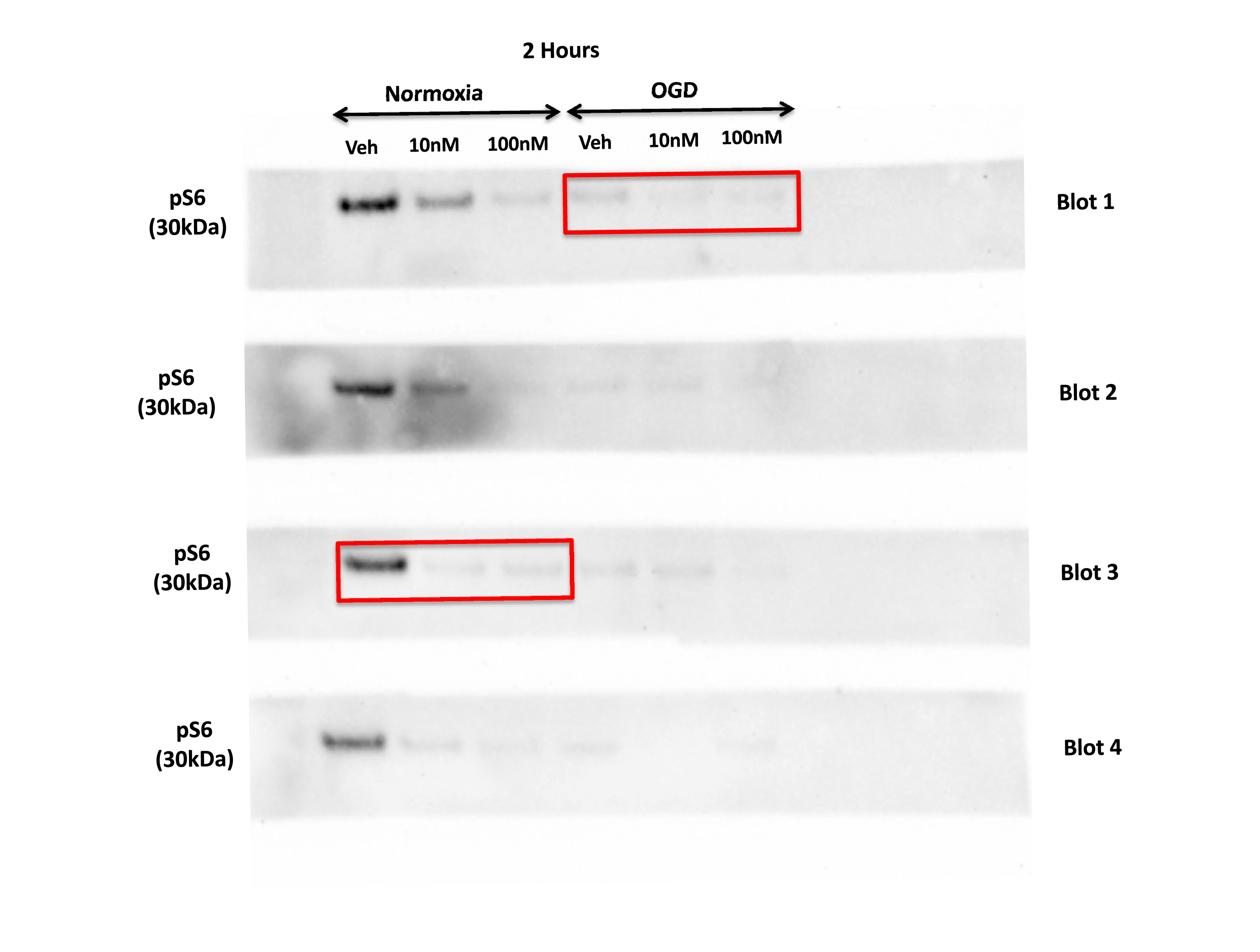

**Supplementary Fig 2** Full unedited blots for Fig. 2 a,d– used for the analysis of Phospho-S6 (pS6) in pericytes exposed to 2 hours of normoxia or Oxygen Glucose Deprivation (OGD). The panels are chemiluminescent images taken using a Biorad ChemiDoc^TM^ MP imaging system, which provides information of the molecular weight/size of the bands (weights depicted to left of blots). The red boxes indicate the bands featured in Fig. 2a,d. The signals of the bands from the original, unprocessed immunoblots were measured using using Biorad Image Lab software v6.0.1. Note, the brightness and contrast of the representative blots in Fig 2 a,d has been enhanced for visualisation purposes only.

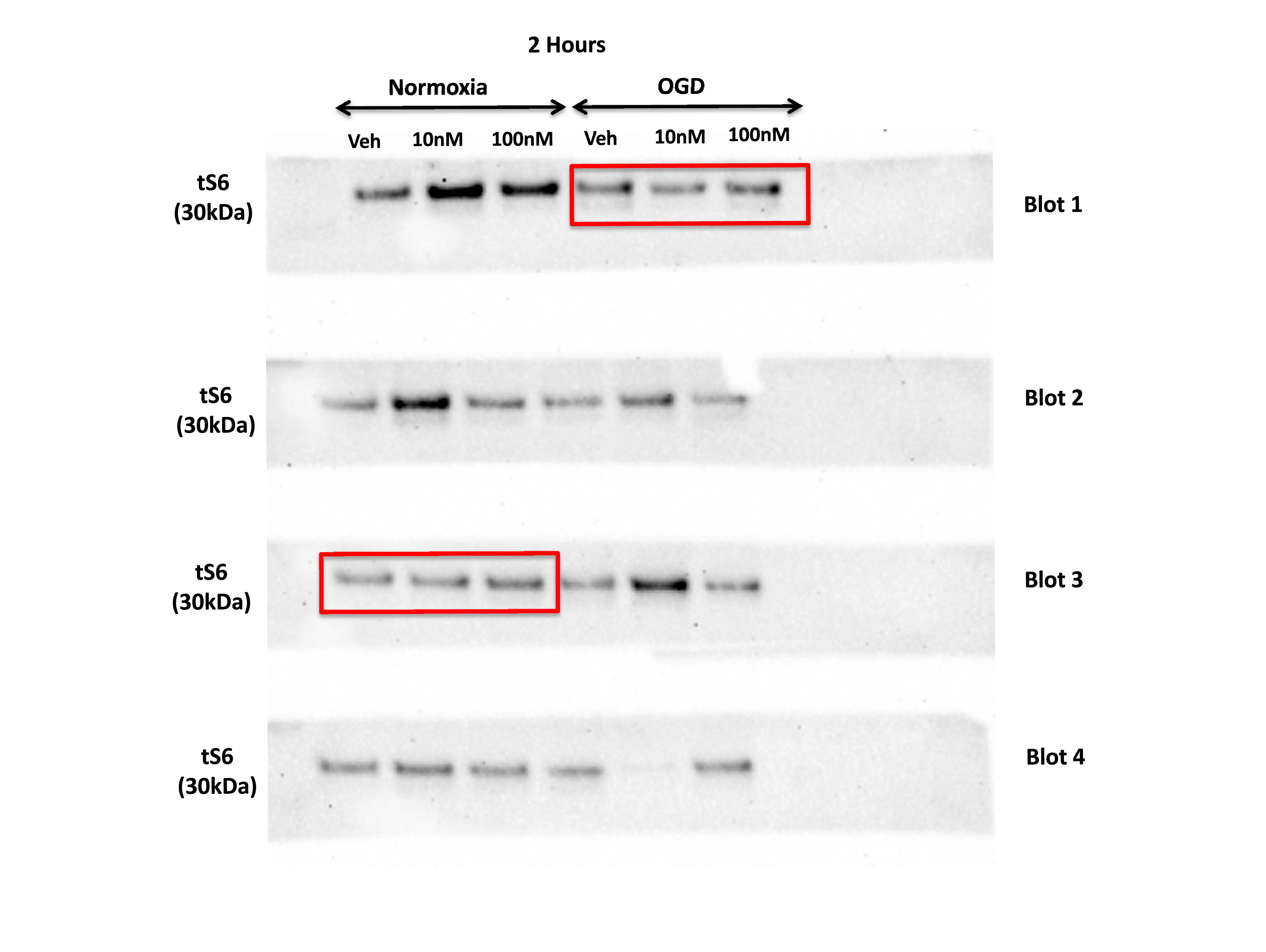

**Supplementary Fig 3** Full unedited blots for Fig 2 a,d– used for the analysis of Total-S6 (tS6) in pericytes exposed to 2 hours of normoxia or Oxygen Glucose Deprivation (OGD). The panels are chemiluminescent images taken using a Biorad ChemiDoc^TM^ MP imaging system, which provides information of the molecular weight/size of the bands (weights depicted to left of blots). The red boxes indicate the bands featured in Fig 2 a,d. The signals of the bands from the original, unprocessed immunoblots were measured using using Biorad Image Lab software v6.0.1. Note, the brightness and contrast of the representative blots in Fig 2 a,d has been enhanced for visualisation purposes only.

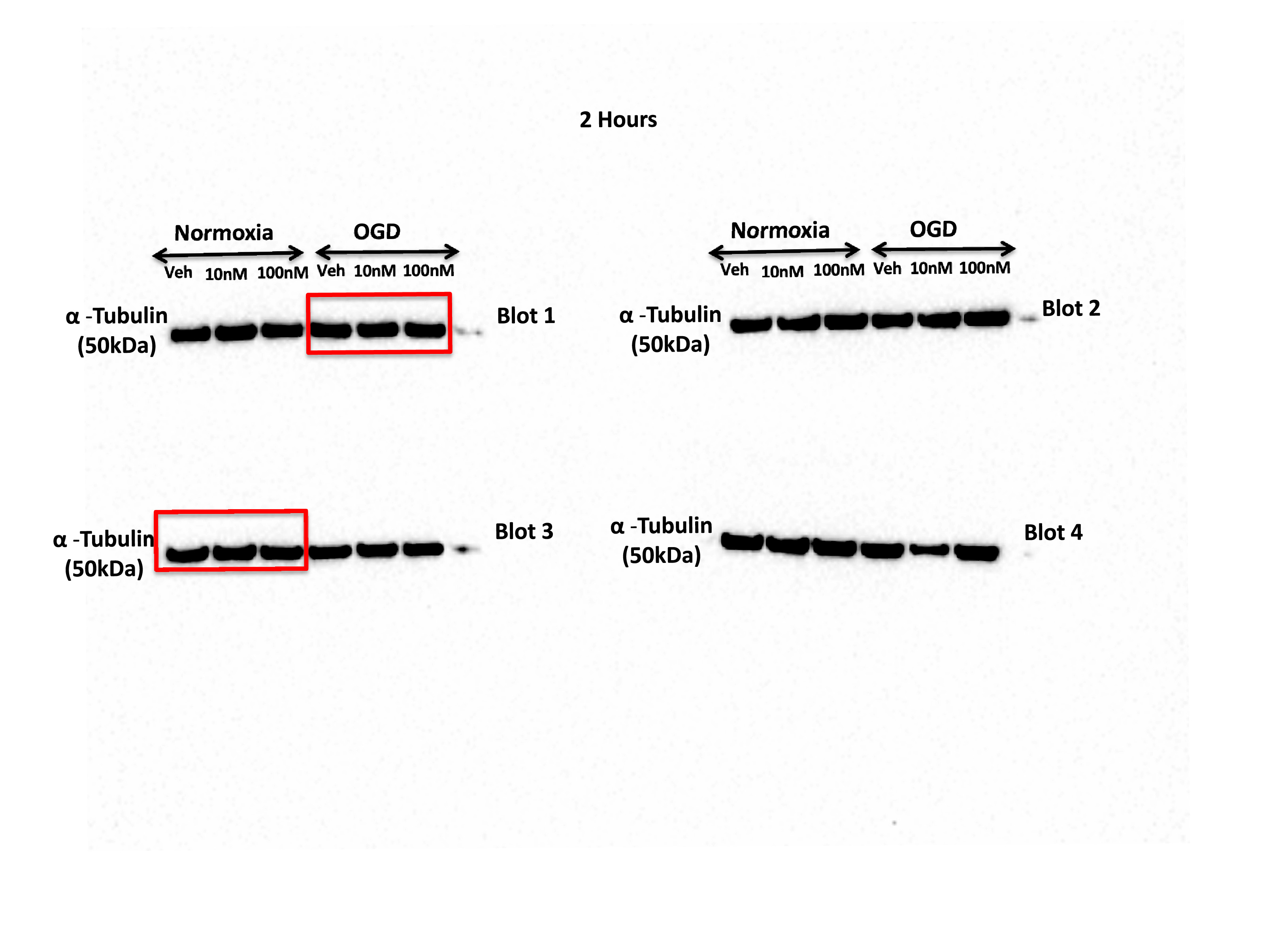

**Supplementary Fig 4** Full unedited blots for Fig 2 a,d– used for the analysis of alpha-tubulin in pericytes exposed to 2 hours of normoxia or Oxygen Glucose Deprivation (OGD). The panels are chemiluminescent images taken using a Biorad ChemiDoc^TM^ MP imaging system, which provides information of the molecular weight/size of the bands (weights depicted to left of blots). The red boxes indicate the bands featured in Fig 2 a,d. The signals of the bands from the original, unprocessed immunoblots were measured using using Biorad Image Lab software v6.0.1. Note, the brightness and contrast of the representative blots in Fig 2 a,d has been enhanced for visualisation purposes only.

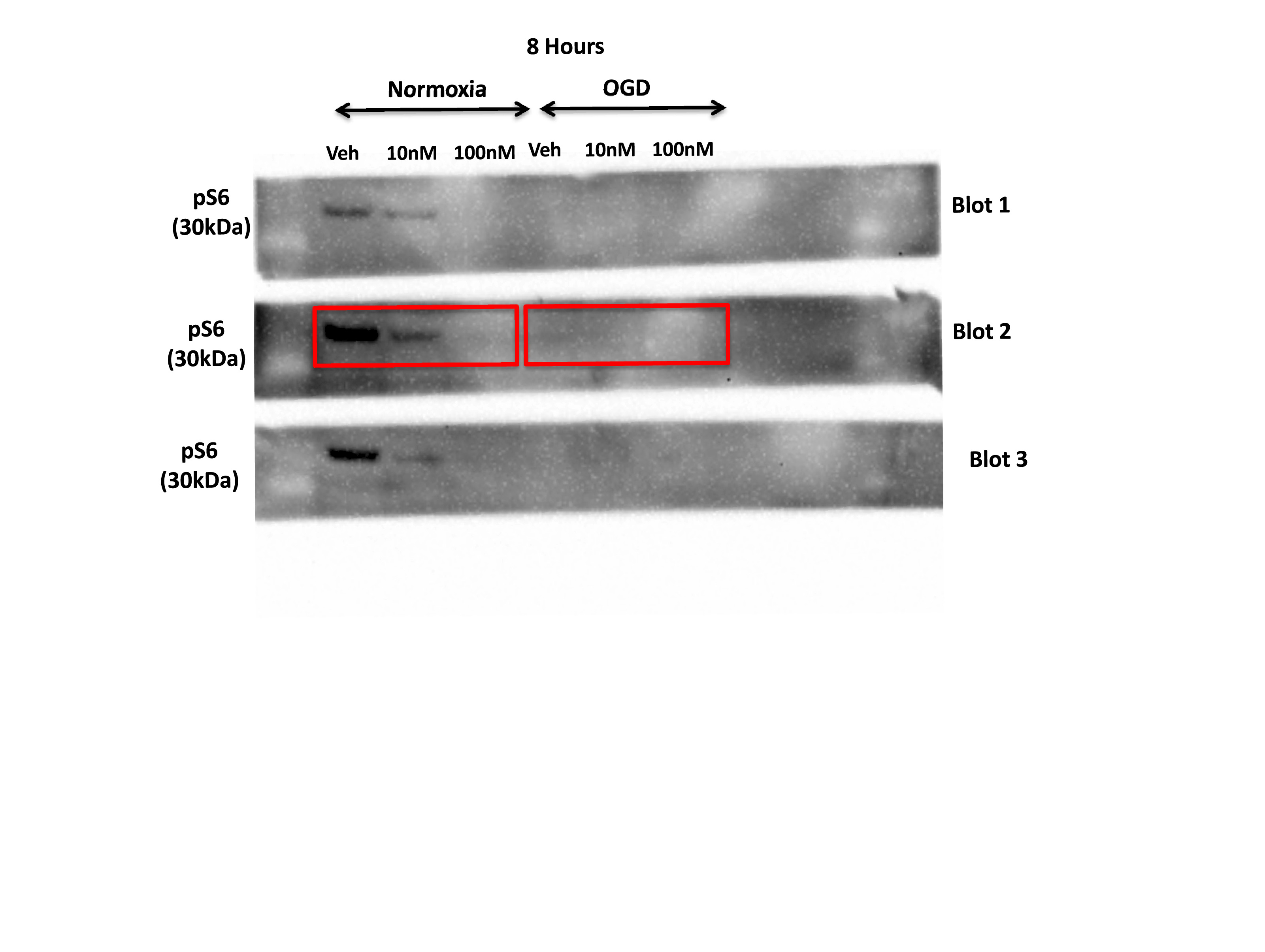

**Supplementary Fig 5** Full unedited blots for Fig. 2 a,d– used for the analysis of Phospho-S6 (pS6) in pericytes exposed to 8 hours of normoxia or Oxygen Glucose Deprivation (OGD). The panels are chemiluminescent images taken using a Biorad ChemiDoc^TM^ MP imaging system, which provides information of the molecular weight/size of the bands (weights depicted to left of blots). The red boxes indicate the bands featured in Fig. 2 a,d. The signals of the bands from the original, unprocessed immunoblots were measured using using Biorad Image Lab software v6.0.1. Note, the brightness and contrast of the representative blots in Fig 2 a,d has been enhanced for visualisation purposes only.

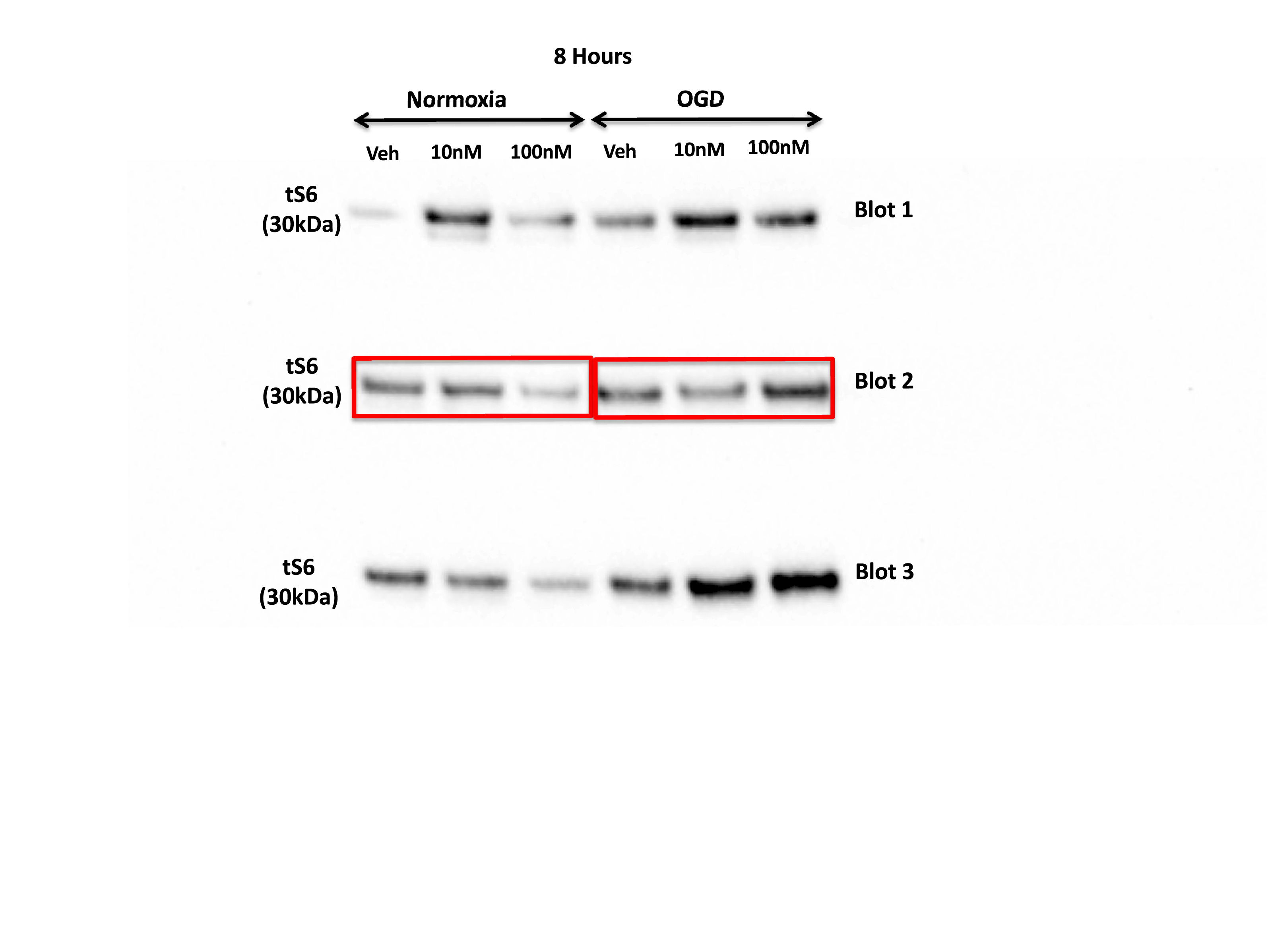

**Supplementary Fig 6** Full unedited blots for Fig 2 a,d– used for the analysis of Total-S6 (tS6) in pericytes exposed to 8 hours of normoxia or Oxygen Glucose Deprivation (OGD). The panels are chemiluminescent images taken using a Biorad ChemiDoc^TM^ MP imaging system, which provides information of the molecular weight/size of the bands (weights depicted to left of blots). The red boxes indicate the bands featured in Fig 2 a,d. The signals of the bands from the original, unprocessed immunoblots were measured using using Biorad Image Lab software v6.0.1. Note, the brightness and contrast of the representative blots in Fig 2 a,d has been enhanced for visualisation purposes only.

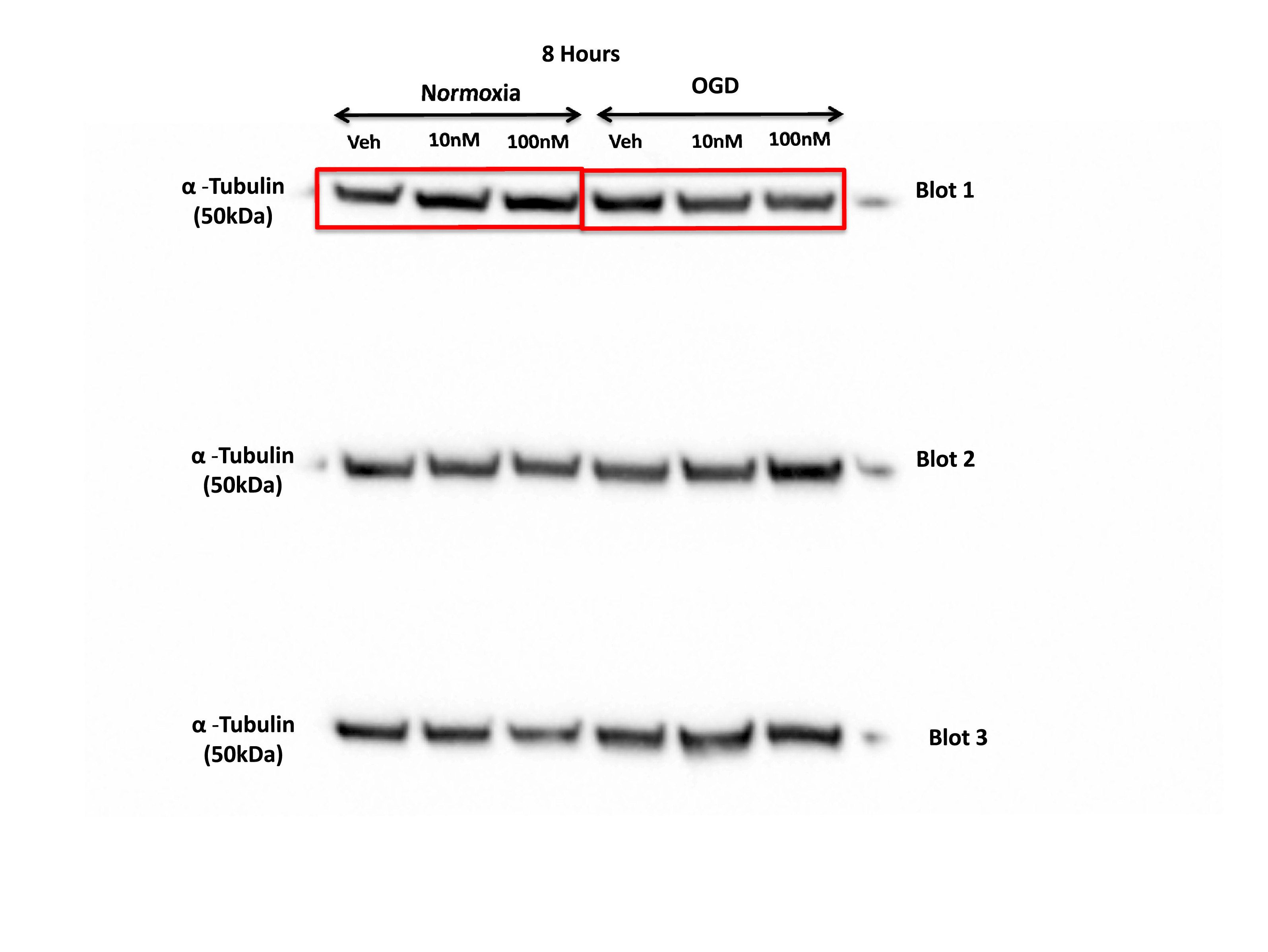

**Supplementary Fig 7** Full unedited blots for Fig 2 a,d– used for the analysis of alpha-tubulin in pericytes exposed to 8 hours of normoxia or Oxygen Glucose Deprivation (OGD). The panels are chemiluminescent images taken using a Biorad ChemiDoc^TM^ MP imaging system, which provides information of the molecular weight/size of the bands (weights depicted to left of blots). The red boxes indicate the bands featured in Fig 2 a,d. The signals of the bands from the original, unprocessed immunoblots were measured using using Biorad Image Lab software v6.0.1. Note, the brightness and contrast of the representative blots in Fig 2 a,d has been enhanced for visualisation purposes only.

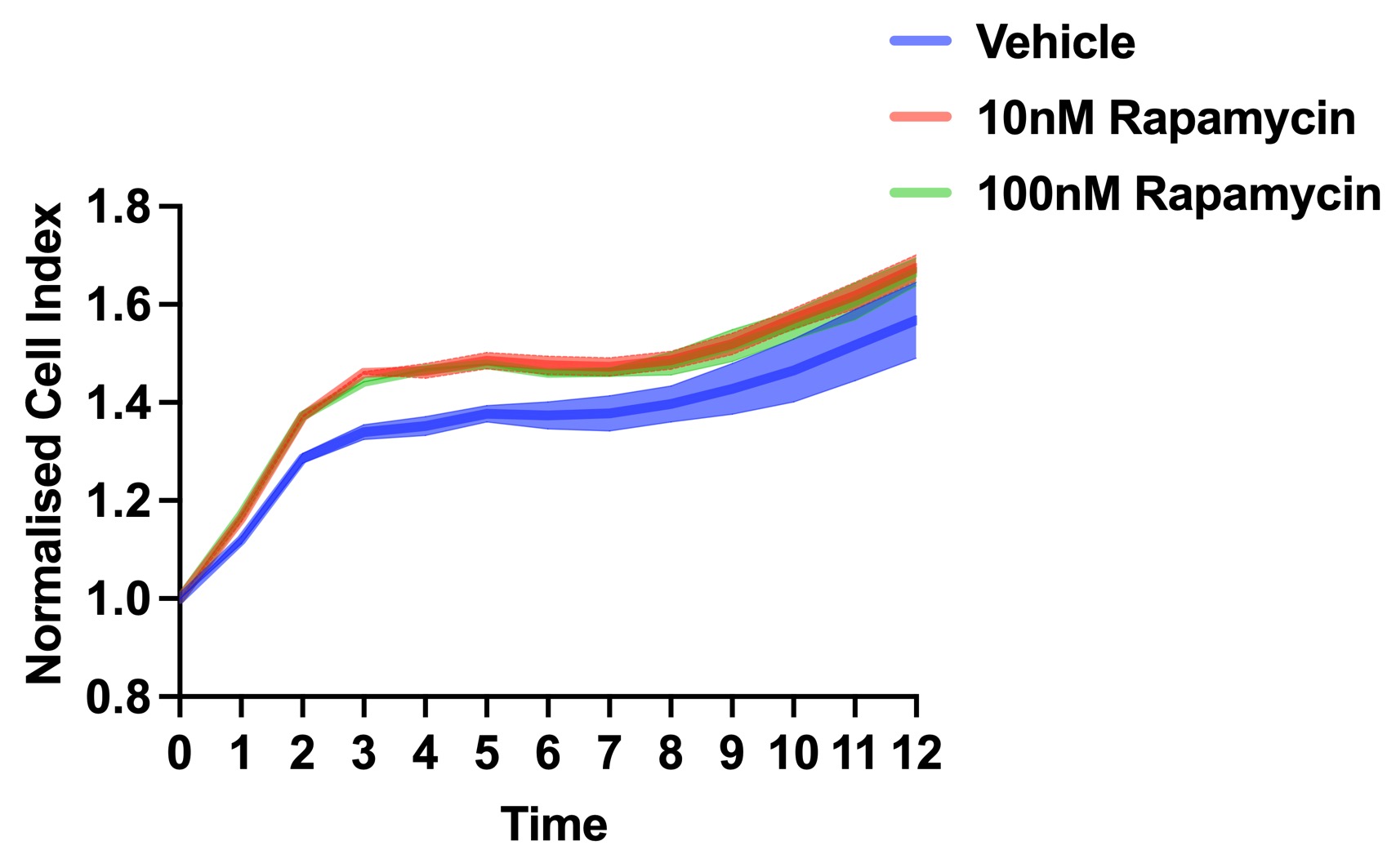

**Supplementary Fig 8** Pericyte contractility during Normoxia. Average normalized cell index for vehicle (black), 10nM rapamycin (light blue) and 100nM rapamycin (dark blue) treated cells during 12 h of normoxia. n = 2 per treatment group.

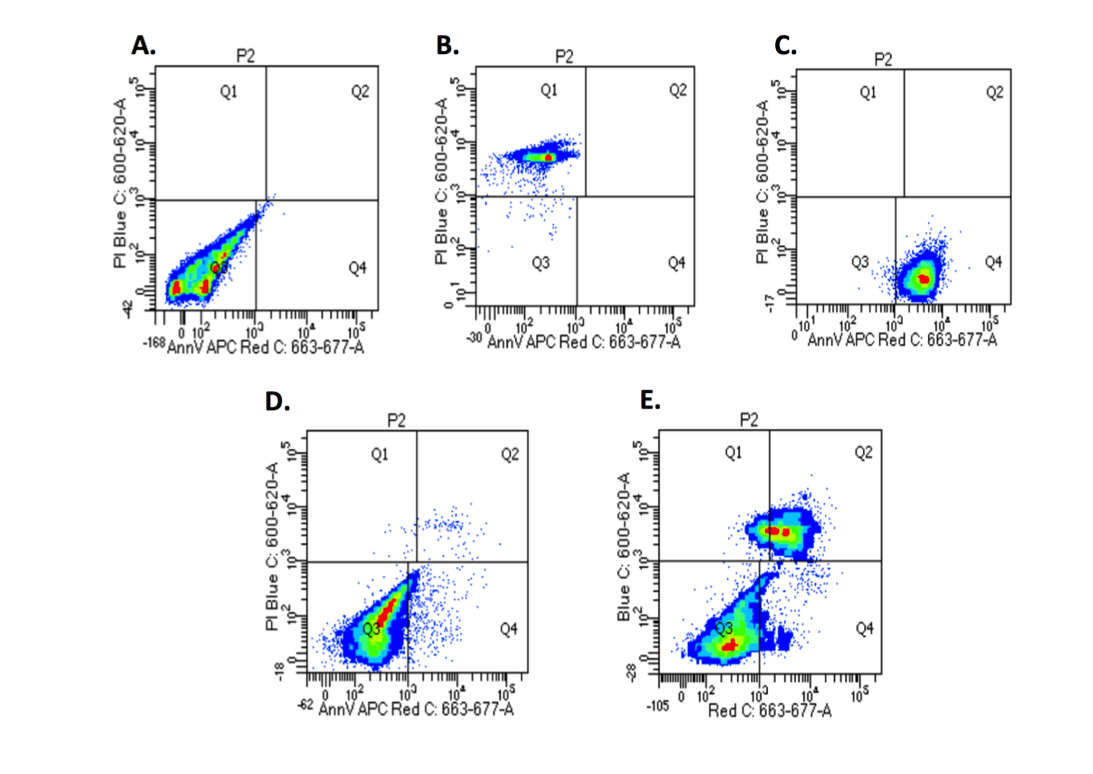

**Supplementary Fig 9** Flow cytometry heat maps. A. Unstained control cells. B. Propodium Iodide (PI) control cells. C. Annexin V (AV) control cells. D. Vehicle Normoxia fully stained cells. E. Vehicle Oxygen Glucose Deprivation (OGD) fully stained cells. Note the increase in the number of cells in quadrant 2 (Q2) indicative of PI-positive and AV-positive cells.

 
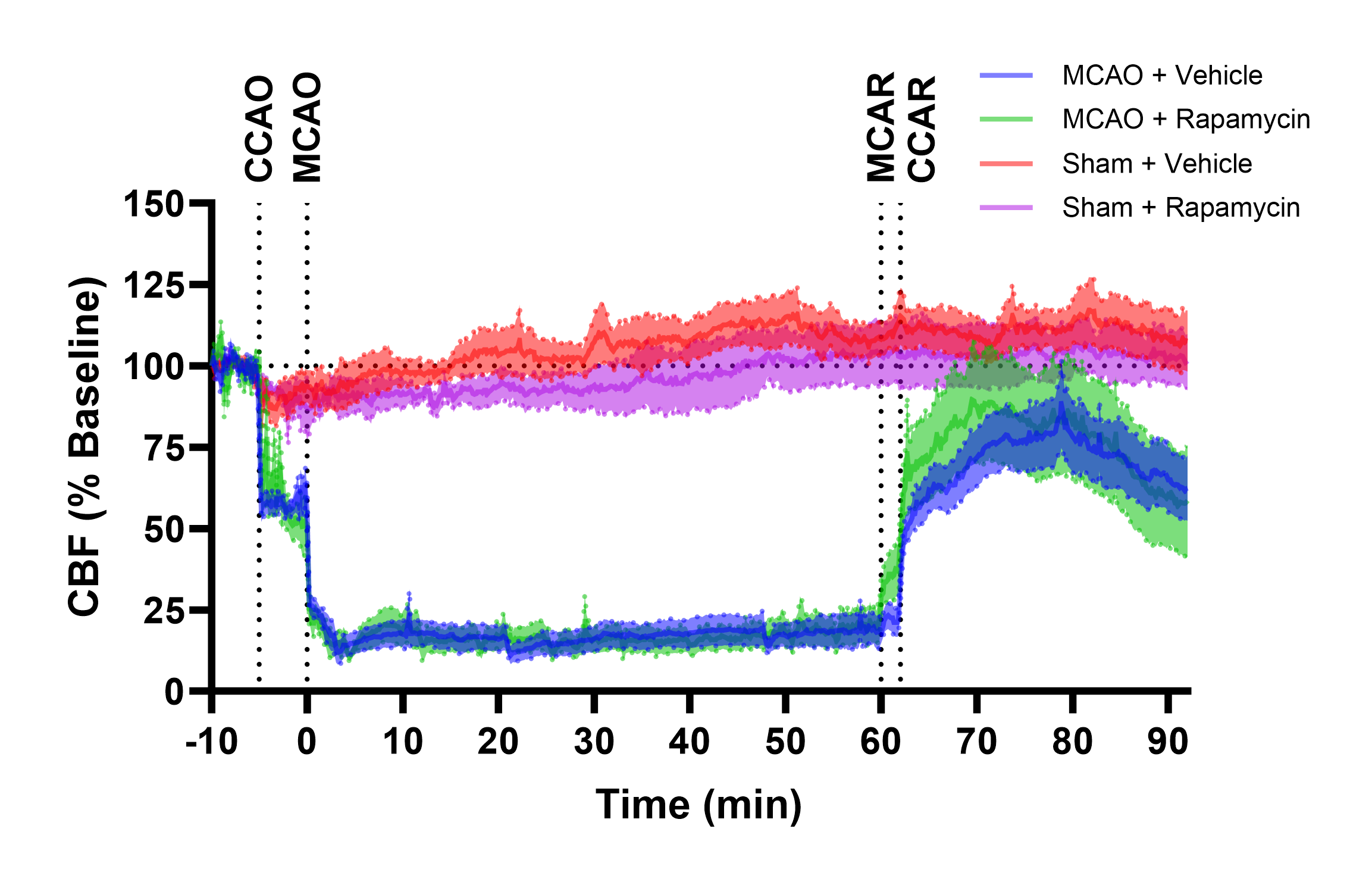
**Supplementary Fig 10** Rapamycin does not affect cerebral blood flow throughout the MCAO and reperfusion periods in the upper somatosensory cortex. Laser-Doppler trace representing cerebral blood flow for NG2-DsRed mice undergoing right-side middle cerebral artery occlusion (MCAO) surgery. Dark lines represent mean CBF and shaded area represents SEM for each treatment group. Number of individual mice analysed for each group: MCAO + vehicle (blue), N=7; MCAO + rapamycin (green), N=8; sham + vehicle (red), N=5; sham + rapamycin (magenta), N=4. Y-axis represents cerebral blood flow (CBF) normalised to a 5-minute section of baseline. X-axis represents time (minutes) normalised to the commencement of MCAO. Horizontal dotted line represents baseline CBF (100%). Vertical dotted lines and corresponding text represent the following: CCAO = time when common carotid artery occlusion occurred; MCAO = time when middle cerebral artery occlusion occurred; MCAR = time when middle cerebral artery recanalisation occurred; CCAR = time when common carotid artery recanalisation occurred.

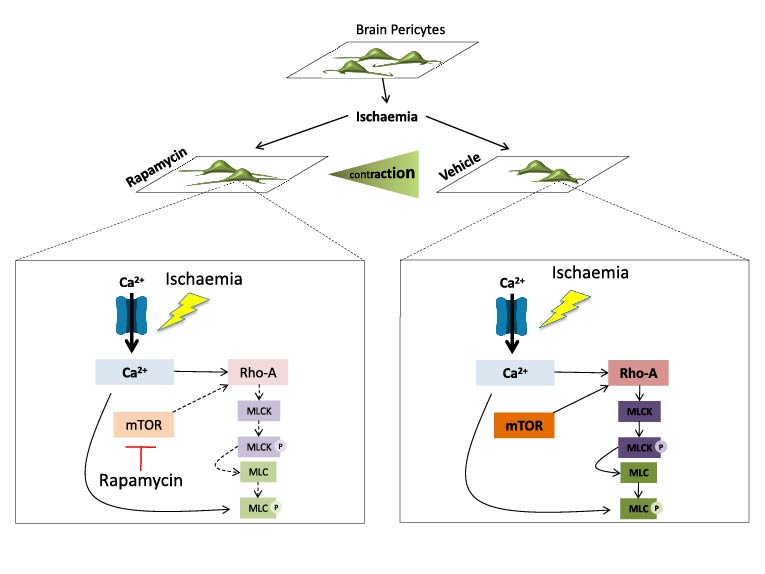

**Supplementary Fig 11** Proposed mechanism of rapamycin on brain pericyte contractility. Calcium and contractile pathways downstream of ischemia leading to pericyte contraction in the absence of rapamycin treatment (right). Rapamycin reduces pericyte contraction to ischemia by reducing mammalian target of rapamycin (mTOR) activity and downstream Ras homolog gene family, member A (RhoA) activity (left). This most likely leads to reduced phosphorylation of the myosin light chain kinase (MLCK) and the MLC. This then reduces sensitivity of the contractile apparatus of the pericyte to the elevated calcium levels during ischemia thus reducing pericyte contraction.
